# Building dynamical models of multi-step state transitions from single cell gene expression trajectories

**DOI:** 10.64898/2025.12.08.693064

**Authors:** Yukai You, Cristian Caranica, Guohao Dai, Mingyang Lu

## Abstract

Multi-step cell state transitions occur across biological processes, such as development and disease progression, yet the underlying gene regulation remains unclear. We introduce NetDes, a computational systems-biology method that infers core transcription factor (TF) regulatory networks and builds ODE-based dynamical models from single-cell gene expression trajectories. In benchmarks on synthetic trajectories with decoys and BEELINE scRNA-seq datasets, NetDes identifies regulatory interactions competitively with existing methods, and reconstructs a simulated cell-fate circuit. We applied it to time-series scRNA-seq data of iPSC-to-definitive-endoderm differentiation, epithelial-mesenchymal transition, erythropoiesis, and dendritic cell differentiation. NetDes has advantages over existing approaches in reconstructing a minimal network with a single model that reproduces observed expression dynamics and captures sequential state transitions. Network simulations predict TFs and their combinations driving each transition, recovering known master regulators, while network coarse-graining reveals the circuit logic of iPSC-to-DE differentiation. NetDes provides a general framework for mechanistic modeling of complex cell state transitions.

## Introduction

Cell state transitions play a crucial role in many biological processes, such as embryonic development(1–3), tissue regeneration(4,5), and disease progression(6,7). Technological advances in genomics(8–10) and live-cell imaging(11) have revealed that cell state transitions often proceed gradually through multiple steps of intermediate states, in contrast to binary state switches. For example, cancer cells can undergo a complete epithelial-mesenchymal transition (EMT) by transiting through hybrid states(12). During the development of hematopoietic stem cells (HSCs), progenitor cells pass through pre-HSC stages before becoming fully functional(13). In mouse embryonic stem cell (mESC) differentiation, the formative state has been identified as an intermediate state between the naive and primed states(3). Elucidating the regulatory principles underlying multi-step state transitions remains a major question in systems biology.

Single-cell RNA sequencing (scRNA-seq) has become a widely used technology for studying cell state transitions, as it enables genome-wide gene expression profiling of individual cells undergoing these transitions. With advances in computational methods, researchers can now map dynamical changes in cell states from scRNA-seq data. For example, pseudotime inference(14,15) and lineage tracing(9) approaches allow cells to be ordered along continuous developmental trajectories, while methods related to RNA velocity(16–18) enable the prediction of single-cell vector fields to represent gene expression dynamics. Building on these dynamics, fate-mapping methods assign each cell a probability of reaching each terminal state(19). However, while these approaches help to describe state transitions, it is still challenging to elucidate the gene regulation of these transitions and accurately predict key regulators capable of driving specific transitions.

Many computational methods have been developed to infer gene regulatory networks (GRNs) from scRNA-seq data, using statistical inference(20–23), machine learning(24–28), or dynamical systems approaches(16,20,29–34). Despite these advances, most existing methods have focused on reconstruction of large networks for individual cell types or discrete cell states, rather than modeling regulatory systems that drive continuous multi-step transitions. For example, scMTNI employs a multi-task learning framework to infer GRNs along a defined lineage, where discrete GRNs are built for individual cell states(24). LINGER uses a neural-network-based method to reconstruct either a cell population GRN, cell-type-specific GRNs or cell-level GRNs(26); however, the method was not designed to construct a GRN capturing state transition dynamics. Furthermore, inferring causal regulatory interactions directly from statistical associations proves to be difficult(35–37), especially when analyzing noisy single-cell datasets that include technical noise such as dropout effects(38) and biological noise derived from stochastic gene expression(39).

In our view, these limitations could be alleviated by the following strategy that integrates top-down and bottom-up systems biology(40). <u>First</u>, a relatively small set of genes, known as transcription factors (TFs), largely regulate the transcription of the other genes, including other TFs, although exceptions do exist. The regulatory interactions among a set of core TFs may form functional gene circuits that perform specific cellular functions, such as gene expression oscillators in the cell cycle(41) and multistable switches governing cell fate determination(42). Thus, we aim to model a core set of TFs and the regulatory interactions among them and evaluate how effectively such core TF networks can capture the observed gene expression dynamics during cell state transitions. <u>Second</u>, to improve the mechanistic inference from noisy single cell data, we incorporate literature-based candidate interactions and denoise the data by smoothing gene expression trajectories along pseudotime, which generates much cleaner inputs for constructing minimal ordinary differential equation (ODE)-based models of a GRN. To robustly identify regulatory signs and parameters from noisy data, we introduce a four-stage optimization procedure with sign-combination sampling, followed by a reducibility index to build a minimal GRN model without sacrificing fit. <u>Finally</u>, based on the optimized mathematical models, subsequent systems biology analyses, such as network simulations of extrinsic signaling and network coarse-graining(43,44), could uncover the dynamical properties of the GRNs. These mechanistic models allow cell state transitions to be interpreted using the Waddington landscape(45,46), a conceptual framework from dynamical systems theory that represents cell states as attractors and state transitions as movement along the landscape defined by the GRN. Such insights into network dynamics and the underlying landscape are often difficult to obtain using conventional network inference methods alone. In the following, we will present the computational method and demonstrate its utility through benchmarking on both synthetic datasets and experimental scRNA-seq data.

## Materials and Methods

### Methodology Overview

We introduce NetDes (<u>Net</u>work inference and optimization using <u>D</u>ynamical <u>e</u>quation <u>s</u>imulations), a computational method for constructing core TF regulatory networks that drive multi-step cell state transitions and building dynamical models using gene expression time trajectories. The workflow of NetDes is outlined in **Fig.1**. First, from time-series scRNA-seq data (the leftmost panel in **Fig.1A**), NetDes infers pseudotime with a higher time resolution. Second, for each gene, the gene expression time trajectory is fitted and smoothed along inferred pseudotime (2^nd^ panel from the left in **Fig.1A**). This step is achieved by applying PseudotimeDE(47), which models each cell’s gene expression as sampled from a negative binomial distribution, where the mean expression along pseudotime is fitted by a spline curve. Third, these genes are clustered according to the shapes of the gene expression time trajectories (3^rd^ panel from the left in **Fig.1A**). Fourth, each context-specific TF is identified whose target genes are enriched in a trajectory cluster (the rightmost panel in **Fig.1A**). This enrichment analysis is performed by assuming that genes with similar time trajectories are likely regulated by the same TF. Fifth, NetDes constructs an initial GRN of the enriched TFs by connecting the genes with regulatory links according to the TF-target databases in TRRUST(48) and the TF binding motifs identified using RcisTarget(49) (1^st^ and 2^nd^ panels in **Fig.1B**). At this stage, the activating/inhibiting nature of regulatory interactions is yet to be determined. Sixth, nonlinear ODEs are fitted against gene expression time trajectories, allowing to trim unnecessary interactions, as well as constructing an ODE-based dynamical model of the optimized GRN (3^rd^ and 4^th^ panels in **Fig.1B**). The established dynamical model also specifies whether a regulatory link is excitatory or inhibitory. Finally, using the established dynamic models, we can perform dynamical systems modeling, including gene expression dynamics simulations for the GRN driven by extrinsic signaling, ensemble-based modeling of state distribution from GRN topology, and network coarse-graining (**Fig.1C**). While conventional GRN inference methods output ranked regulatory edges, NetDes provides a mechanistic model of core TFs, together with predictions of top driver genes and their combinations in driving desired state transitions. The step-by-step procedure for NetDes is detailed in the following section.

**Fig. 1.**
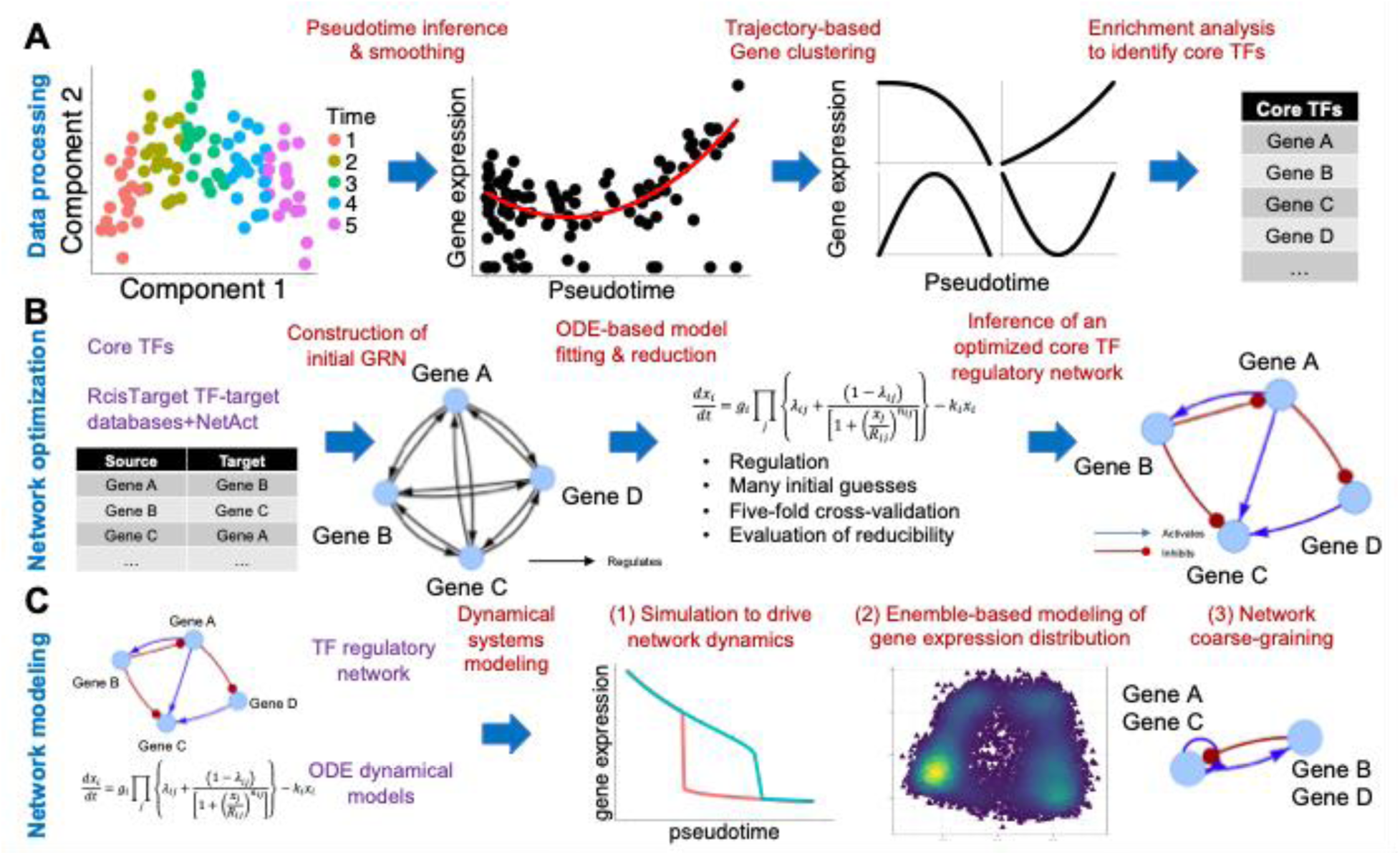
Overview of NetDes for inferring GRN and the associated dynamical model using scRNA-seq data. **(A)** Data processing. From time-series scRNA-seq data, we first infer pseudotime with a higher time resolution. We then apply PseudotimeDE to fit a smooth time trajectory for each gene. Genes are then clustered according to the shapes of their time trajectories. From the gene clusters, we identify core transcription factors (TFs) whose target genes are enriched in the time trajectory clusters. **(B)** Network optimization. An initial gene regulatory network (GRN) is built by connecting the inferred core TFs with regulatory interactions from literature-based TF-target databases. From the initial GRN, we optimize a minimal GRN and the associated nonlinear ODE-based dynamical model to match gene expression time trajectories. **(C)** Network modeling. With the established GRN model, we can perform (1) simulations to drive cell state transitions with extrinsic signals to network genes; (2) ensemble-based modeling to explore robust network states; (3) network coarse-graining to infer core gene circuits.

A major component of NetDes is a robust algorithm for fitting and optimizing a GRN and the associated nonlinear ODE model (the sixth step mentioned above). For every network gene, the kinetic parameters for the gene’s ODE are fitted according to the regulators’ and target’s smoothed gene expression trajectories by a four-stage model optimization (see **Supplementary Methods**, **Fig.S1** for method details, and **Fig.S2** for an example of optimization results). To prevent the nonlinear fitting from being trapped in local minima, NetDes performs model fitting over multiple rounds to sample different combinations of regulators’ activating and inhibiting types. To ensure robust model fitting, NetDes incorporates regularization terms, five-fold cross-validation (CV), and a model reduction guided by a reducibility index that measures the relative loss in fit accuracy after reduction. Overall, these strategies ensure a reliable dynamical model generation of minimal GRNs capturing multi-step state transitions.

### Single-cell RNA-seq datasets

In this study, we analyzed four time-series scRNA-seq datasets representing different cases of cell-state transitions. The first dataset comes from an experiment profiling human induced pluripotent stem cell (iPSC) differentiation into definitive endoderm (DE) induced by a cytokine and small-molecule cocktail (50), for 125 donors at four time points (Day 0, Day 1, Day 2, and Day 3). The second dataset profiles A549 lung adenocarcinoma cells undergoing TGF-β1–induced epithelial-mesenchymal transition (EMT) (51), sampled at Day 0, Hour 8, Day 1, Day 3, and Day 7; cells from the washout conditions were excluded prior to analysis. The third dataset consists of profiles from mouse bone-marrow myeloid progenitors (52); although the original dataset includes multiple lineages, we retained only cells from the erythroid lineage to model erythropoiesis (ERY). The fourth dataset is derived from a temporal map of human in vitro myelopoiesis from iPSC-derived cells (53); we retained cells from the Day-31 embryoid-body myeloid stage together with the subsequent DC-phase time points (Day 31+1, Day 31+4 and Day 31+7), restricted to the monocyte–DC-precursor and cDC2 populations in the G1 cell-cycle phase. All four datasets were preprocessed and subsequently analyzed using NetDes for gene regulatory network modeling, as described below.

### Trajectory inference, smoothing, and clustering

#### Pseudotime inference

Time-series scRNA-seq datasets were obtained as counts per million (CPM) for the iPSC-to-DE differentiation dataset and as raw UMI counts for the EMT, ERY and DC datasets. All four datasets were processed with the same pipeline: counts were processed by sctransform(54) for normalization. The normalized data was then *log*_2_ -transformed (*log*_2_(normalized_data + 1) ) and standardized (referred to as the processed data below). We conducted principal component analysis (PCA) on the top 500 biological variable genes (obtained by scran(55) using the CPM data) from the processed data and subsequently inferred pseudotime by Slingshot(56), using the first five principal components (PCs) for the iPSC-to-DE and DC datasets, the first three PCs for the EMT dataset and the first two PCs for the ERY dataset. Pseudotime inference allows us to characterize gene expression trajectories for NetDes to construct and optimize GRN dynamic models.

#### Trajectory smoothing

Using the pseudotime value and single cell gene expression for each cell, we applied PseudotimeDE(47) to infer a smoothed time trajectory of gene expression for each of the top 5000 highly variable genes (HVGs) identified by scran. Here, PseudotimeDE assumes that each gene in each cell follows a negative binomial distribution and average gene expression along pseudotime is fitted by a spline curve with four knots (see **Fig.S3** for examples of fitted trajectories for selected TFs). Each smoothed gene expression trajectory was then standardized to have zero mean and unit variance.

#### Trajectory-shape-based gene clustering

These top 5000 HVGs were clustered according to the smoothed gene expression time dynamics. We devised a new approach for clustering because of low performance of several existing algorithms, such as K-means, K-medoids and Hierarchical clustering, in clustering time trajectories (**Fig.S4**). First, we excluded genes whose processed gene expression data had a standard deviation below a threshold level (0.25). Second, for the rest of the genes, we classified time trajectories according to the increasing and decreasing patterns of gene expression dynamics. To achieve this, for each smoothed time trajectory, we first determined all possible turning points to partition the trajectory into several small segments. For any segment with maximum gene expression changes smaller than 0.2, we excluded the turning points of the segment (*i.e.*, one turning point for the first or the last segment, and two turning points for the rest). We can exclude these turning points here because the preceding and following segments should have the same increasing and decreasing trends. Finally, we grouped trajectories into gene clusters according to the increasing and decreasing trends of the curve segments delimited by the remaining turning points, as illustrated in **Fig.S4**.

### Construction of an initial TF regulatory network

To obtain key transcription factors (TFs) driving these gene expression dynamics, we hypothesized that genes with similar gene expression trajectories are likely regulated by common TFs. Thus, we performed enrichment analysis on each gene cluster using a literature-based TF-target gene set database, derived from NetAct(57), which compiles databases from TRRUST(48), RegNetwork(58), TFactS(59) and TRED(60). Here, Fisher’s exact test was performed to evaluate the overlapping between genes in a cluster and known target genes for a TF. Here, TFs with fewer than five target genes in the database were excluded. For the iPSC-to-DE, EMT and DC datasets, the background consisted of all genes present in the corresponding scRNA-seq expression matrix (11231, 13239 and 33538 genes, respectively), and the TFs with adjusted p-value less than 0.1 were selected for building GRNs. For the erythropoiesis dataset, the deposited matrix contains only 3,451 genes, 2,444 of which fall into the clusters; the background was therefore restricted to all the 5,353 target genes for the expressed TFs in the mouse TF-target database from NetAct (mDB), and the adjusted p-value cutoff was selected to be 0.3. To avoid finding false positive TFs due to excessive overlap in target genes, we tried to find TF pairs when overlap occurred between the targets of two TFs (q-value < 10^-40^ in the fisher exact test). Specifically, for the iPSC-to-DE case, we retained LEF1 over TCF7L2 as LEF1 showed higher variance in our dataset, whereas TCF7L2 is primarily associated with later gut lineage functions(61). Similarly, between the AP-1 component FOS and JUN, we selected FOS, which displayed greater expression variability during iPSC differentiation(62,63). The only exception is Rara/Rarg in the ERY dataset (q = 2.9 × 10^-62^), in which both were retained, as there is evidence to show they have distinct, non-redundant roles in bone-marrow haematopoiesis(64). Apart from those pairs, no other TF pair was found with significant overlap in the other two datasets.

Once enriched TFs were identified, we constructed an initial TF regulatory network by connecting the TFs according to the regulatory interactions presented in both the literature-based TF-target databases from TRRUST(48) and the RcisTarget(49) database that is derived from cis-regulatory motif prediction. Here, we kept TF-to-TF edges from RcisTarget whose Normalized Enrichment Scores (NES) are above one to include more potential edges in the initial network, with the expectation that unrelated edges will be removed during GRN optimization.

### Optimizing a nonlinear dynamic model of GRN

For each TF in the initial GRN, we optimized a nonlinear ODE model according to the smoothed gene expression time trajectories of itself and its TF regulators (**Fig.S1**). To ensure uniformity in trajectory fitting and reduce the effects of cell distribution variability along pseudotime, we selected *M* = 201 evenly spaced time points along each trajectory. Additionally, we scaled the gene expression values along each trajectory by their maximum value.

The ODEs of the dynamic model were adopted from a common approach in systems-biology modeling of gene regulatory circuits/networks.

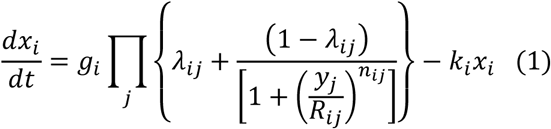

where *x_i_* represents the expression levels of target gene *i*, *y_j_* represents the expression levels of regulator *j*; *g_i_* and *k_i_* are the basal production rate and degradation rate for gene *i* (*k_i_* was fixed to 0.1 for all genes; only the ratio 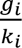 is identifiable from the fit, see **Eq.2**); the product is over all *k_i_* regulators *j* of gene *i* ; *λ_ij_* , *R_ij_* and *n_ij_* are the maximal fold change, Hill threshold, and Hill coefficient for the regulation from regulator *j* to gene *i*(65). Gene *j* activates *i* when *λ_ij_* > 1, and inhibits *i* when *λ_ij_* < 1.

Under the quasi-steady state assumption during the dynamic progress, **Eq.1** leads to

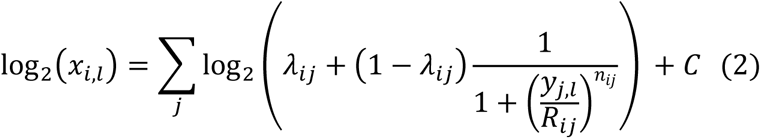

where 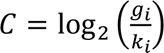. The additional subscription *l* represents the indices of the data points along the time trajectory. To allow unnecessary interactions to be trimmed, we performed nonlinear fitting, for each gene *i*, with a regulation term:

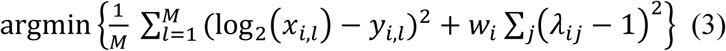

In the fitting, log_2_(*x_i_*_,*l*_) is computed by using **Eq.2**, *y_i_*_,*l*_ for each gene is obtained from the smoothed gene expression trajectories, and *w_i_* is the tuning parameter for the regulation term. The summation is over all *M* = 201 data points. When the optimized *λ_ij_* approaches one, the interaction from genes *j* to *i* is essentially removed from the dynamic model. But instead of directly removing interactions by a LASSO process, we performed four rounds of model fitting (as described in **Supplementary Methods**) by considering all possible subsets of regulators, from which we identified a more robust prediction of regulators. In the ODE models, we also set kᵢ = 0.1 day⁻¹ for all genes, corresponding to slow simulation dynamics that approximate quasi-steady-state behavior during state transitions.

### Model simulations of GRN dynamics driven by extrinsic signals

After the GRNs and the associated ODEs were obtained from model optimization, the gene expression dynamics of the GRNs can be simulated by integrating the ODEs with the 4^th^ Order Runge-Kutta Method. In the simulations, we started from an initial condition where gene expression levels were selected from the initial time point of the smoothed time trajectories and simulated the ODE model until it reaches to a stable steady state. This stable steady state was used as the initial condition for further simulations. To evaluate the dynamical response of the GRN to extrinsic signals, we performed two simulations on the dynamic model by driving the system with the observed gene expression trajectories on one, two or three TFs in the GRN. In the first simulation, the system starts from the initial state and is driven by the gene expression trajectory of the TF (or TFs) along the forward time direction. In the second simulation, the system starts from the final state of the first simulation and is driven by the gene expression trajectory of the TF (or TFs) along the backward time direction. These two simulations allow us to evaluate how effectively the whole system is driven by extrinsic signals targeting selected TF(s), enabling cell state transitions along both forward and reverse directions.

To compare the quality of model fitting and the ability of network genes to drive cell state transitions across datasets on a common scale, we quantified the agreement between a model trajectory (from gene-driving simulation) and the observed trajectory (the reference) by Nash-Sutcliffe efficiency (NSE):

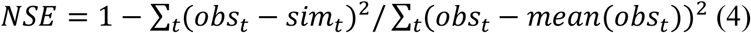

where *obs_t_* is the observed scaled expression at pseudotime point t, *sim_t_* is the model-simulated scaled expression at the same pseudotime point, and mean(*obs_t_*) is the average of *obs_t_* over all t. Both trajectories were min–max scaled to [0,1] and have 201 time points. NSE = 1 indicates perfect fitting, a value of zero indicates fitting no better than predicting the mean, and a negative value indicates fitting worse than the mean.

### Benchmarking design

NetDes was benchmarked against existing GRN inference methods in the following three tests (details in **Supplementary Methods**). The first benchmark tested whether ground-truth regulators could be distinguished from decoy genes (non-regulators) using synthetic trajectories, where the target gene’s dynamics were generated by a model with the regulators and their activating/inhibiting nature pre-defined by construction. Performance was evaluated by Area Under the Receiver Operating Characteristic curve (AUROC) and Area Under the Precision-Recall Curve (AUPRC) against the known regulators.

The second benchmark tested whether cell-type-specific TF-target relations (regarded as the ground truth) could be correctly distinguished from non-specific relations within the combined relation set by inference methods. Here, we tested on human embryonic stem cell (hESC) and hepatocyte (hHep) datasets from the BEELINE collection, with scRNA-seq data used as input and ChIP-seq data used to define cell-type-specific and non-specific TF-target relations. Performance was assessed by the mean AUPRC across target genes. Note that this test only evaluates identification of individual edges, not their collective effects on cell states, which is what NetDes is designed to capture.

The third benchmark tested whether an inferred GRN robustly reproduces the cell states along the observed state transitions using four time-series scRNA-seq datasets (iPSC-to-DE differentiation, EMT, erythropoiesis, and DC differentiation). Here, each inferred GRN was simulated by Random Circuit Perturbation (RACIPE) (65–68), an ensemble-based gene circuit simulation method, to generate a population of gene expression profiles from models with randomly sampled kinetic parameters. These profiles were then evaluated against reference states sampled along the measured state-transition trajectory using weighted mapping percentage, the entropy of the mapped state distribution, and scaled distance, as defined in detail in **Supplementary Methods**. For the iPSC-to-DE differentiation case, inferred GRNs were also assessed for agreement with literature-curated interactions.

Tested GRN inference methods include ppcor, GENIE3, SCODE, DeepSEM, DeepRIG, SINCERITIES and CellOracle, whose usage is described in detail in **Supplementary Methods**. In the third benchmarking test, because NetDes begins from a literature-derived initial GRN, all methods were restricted to this initial GRN before evaluation, except for CellOracle, which uses its own initial GRN. It is also worth noting that NetDes ranks combination of regulators rather than individual edges; to ensure consistent performance evaluation, such rankings were converted into an edge ranking according to each regulator’s appearance among top-ranked combinations. However, such conversion may understate NetDes’s performance.

### GRN Coarse-graining

We applied a coarse-graining method, Sampling Coarse-Grained Circuits (SacoGraci)(44), to construct small gene circuits from the optimal core TF GRN inferred by NetDes. SacoGraci constructed coarse-grained circuits by a Markov Chain Monte Carlo sampling of small gene circuits, whose simulated gene expression states match those from the full GRN. When applying to the iPSC-to-DE differentiation case, we manually categorized genes into three groups according to their RACIPE simulated expression patterns along pseudotime (early, intermediate and late activation). In addition, simulated gene expression profiles from the full GRN were clustered by using hierarchical clustering analysis (three clusters, distances based on signed Pearson correlation, and Ward.D2 linkage). The coarse-grained gene circuit was then modeled to have nodes representing gene clusters and circuit states representing clusters for gene expression profiles. We used the default parameters for SacoGraci and ran the optimization 30 times to obtain the best coarse-grained circuit.

## Results

### NetDes distinguishes true regulators from decoys in synthetic benchmarking

A major challenge in GRN inference is to distinguish true regulators of a target gene from genes whose expression is only associated with it. To test whether NetDes meets this challenge, we performed a comprehensive benchmark on an *in-silico* dataset against several popular network inference methods, *i.e.*, ppcor(22), GENIE3(25) and SCODE(30) (**Fig.2A**). In this benchmark test, we randomly generated ten time trajectories from polynomials (**Eq. S4**) to represent regulators’ expression dynamics (**Fig.S5**). For each case, a subset of randomly selected regulators (serving as ground truth) was assigned to a target gene, and the target gene’s expression dynamics were simulated using **Eq. S5** with a set of randomly sampled kinetic parameters. Each test provides as the input the trajectories of the regulators and target, alongside the trajectories of randomly selected non-regulators (serving as decoys). From a total of 81 tests, we evaluated how various network inference methods can correctly identify the true regulators of the target gene (see **Methods**for details). From the benchmarking, we found NetDes outperforms the other methods in identifying the ground truth regulators according to both AUPRC (leftmost points in the top left panel in **Fig.2B**) and AUROC (**Fig.S6**). NetDes’s performance was robust in the benchmark test when noise of different levels was added to the gene expression time trajectories (top left panels in **Fig.2B** and **Fig.S6**). When evaluating the outcomes of the inference for activators and inhibitors separately, we found all methods performed slightly worse for predicting inhibitors, whereas NetDes still performed better (middle and right panels in **Fig.2B** and **Fig.S6**). Moreover, the performances of various methods became worse when testing on cases with more decoy genes, as these tests are likely more challenging (bottom panels in **Fig.2B** and **Fig.S6**).

**Fig. 2.**
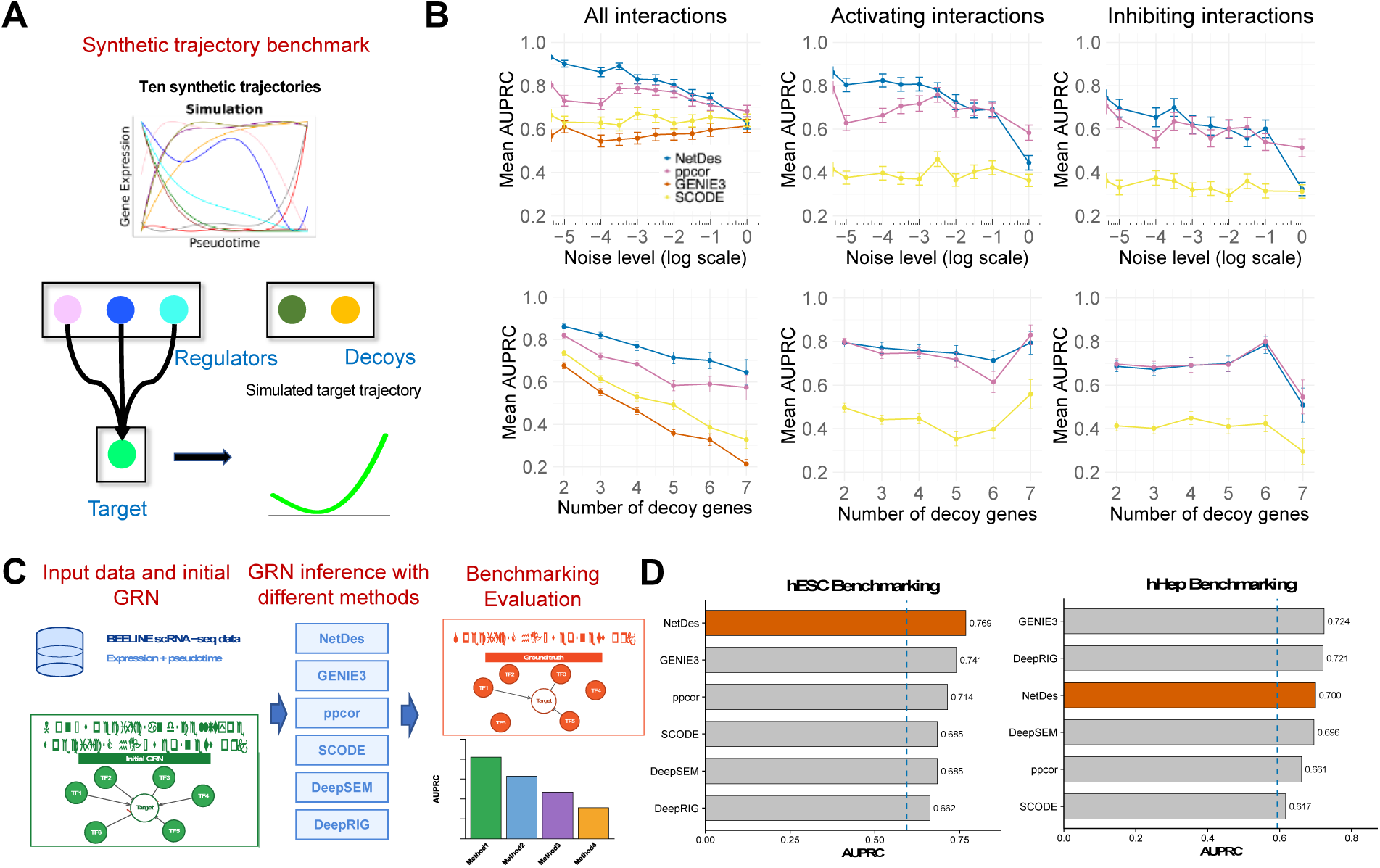
Benchmarking network inference using synthetic trajectories with decoys and experimental scRNA-seq data. **(A)** Illustration of the process used to generate the synthetic dataset for benchmarking. First, different gene expression time trajectories are generated for ten genes. Second, for each test case, a subset of these genes is randomly selected as regulators. Their gene expression trajectories are then used to simulate the trajectory of a target gene according to an ODE model with randomly selected kinetic parameters. Third, additional genes from the remaining set are chosen as decoys. Finally, in each test, the time trajectories of true regulators, target gene, and decoys are provided as input to network inference methods. The goal is to evaluate whether a network inference method can effectively distinguish true regulators from decoys. **(B)** Benchmarking results for the synthetic dataset. Top panels show the AUPRC of different network inference methods applied to simulated gene expression data under different noise levels. Bottom panels show the mean AUPRC of various methods when applied to cases with different numbers of decoy genes. The leftmost, middle and rightmost columns show the benchmark results for all interactions (i.e., both activating and inhibitory), activating interactions, and inhibitory interactions, respectively. **(C)** Illustration of the benchmark design on experimental scRNA-seq datasets from the BEELINE collection. For each dataset, the processed expression matrix together with pseudotime ordering is provided to each inference method, which returns a ranked list of inferred regulatory edges. The initial GRN is defined by the union of a non-specific ChIP-seq network and a cell-type-specific ChIP-seq network, while the cell-type–specific ChIP-seq network serves as the ground truth. Only the inferred edges already existing in the initial GRN were considered for evaluation, and performance is quantified by the average AUPRC across target genes. **(D)** Benchmarking results on the hESC (top) and hHep (bottom) datasets. Bars show the average AUPRC for NetDes (highlighted in orange), GENIE3, ppcor, SCODE, DeepSEM and DeepRIG. The dashed vertical line indicates the expected AUPRC of a random predictor, i.e. the average fraction of true regulators among the candidate regulators across target genes.

To further evaluate the performance of NetDes, we benchmarked it using two single cell RNA-seq datasets from BEELINE collection(35), i.e., human embryonic stem cells (hESC) and hepatocytes (hHep) against network inference methods GENIE3 (23), ppcor (20), SCODE (26), DeepSEM(69) and DeepRIG(70). In this benchmark test, each method was provided with the processed expression matrix and pseudotime ordering from BEELINE, and returned a ranked list of regulatory edges. We defined the initial GRN as the union of a non-specific ChIP-seq network and a cell-type-specific ChIP-seq network, and used the cell-type-specific network as the ground truth (**Fig.2C**). The performance was evaluated by the mean AUPRC across target genes. We found NetDes achieved the highest mean AUPRC on hESC (0.769, compared with 0.741 for GENIE3, 0.714 for ppcor, and 0.63 for the random-predictor baseline), and remained among the best-performing methods on hHep (0.700, compared with 0.59 for the random-predictor baseline), close to GENIE3 (0.724) and DeepRIG (0.721) (**Fig.2D**). Together, the benchmarking results demonstrate that NetDes can identify true regulatory interactions competitively with existing network inference methods on both synthetic trajectories and experimental scRNA-seq data.

### NetDes reconstructs a simulated ICM-trophectoderm switch

Beyond identifying individual regulatory interactions, a key test is whether NetDes can reconstruct, from simulated gene expression data, a dynamical model of a biological gene circuit driving specific cell state transitions. To address that, we investigated into a three-node circuit, consisting of Oct4, Cdx2 and Esrrb, that governs earlier embryonic development in mouse. Oct4 maintains the inner cell mass (ICM) identity, whereas Cdx2 drives extraembryonic trophectoderm (TE) commitment. Around the early blastocyst, ICM cells retain the plasticity to reversibly switch to the TE fate but can become committed to the TE lineage through the activation of the FGF/MAPK signaling pathway(71).

Here, we utilized this biological circuit to generate simulated gene expression trajectories and evaluated how well NetDes can reconstruct the circuit and recover its cell state transition properties through a dynamical model. To establish the ground-truth gene circuit (top left panel in **Fig.S7**), we conducted a manual literature survey to identify relevant gene regulatory interactions as follows. It has been shown that Esrrb activates Oct4 transcription in embryonic stem cells(72), and Cdx2 represses Esrrb transcription in trophoblast stem cells according to ChIP-seq evidence(73). Oct4 and Cdx2 form a toggle switch through mutual inhibition(74), while the incorporation of Esrrb adds a coherent feedforward loop. We also included FGF as an input signaling node that drives the circuit directly through Cdx2(71).

To generate the simulated dataset for gene circuit reconstruction, we establish an ODE model which allows two stable steady states corresponding to the ICM and TE states, respectively. The detailed ODEs and model parameters are provided in **Table.S2**. We then simulated the expression dynamics of Oct4, Cdx2 and Esrrb by driving the circuit from the ICM state to the TE state through an increasing FGF signal (bottom left panel in **Fig.S7**). For network modeling tests, we applied NetDes twice on two initial networks: the first was a fully connected network, while the second lacked the interaction from Oct4 to Esrrb, making it closer to the ground truth. In both cases, the simulated gene expression trajectories of all genes were inputted to NetDes.

From circuit reconstruction, NetDes recovered the ground truth circuit when starting from the second initial GRN, whereas it inferred an additional activation from Oct4 to Esrrb when starting from the first initial GRN. When simulating the optimized GRN models, we found, in both cases,

FGF successfully drove the system in both directions. However, only in the second case could the circuit dynamics be driven backward by directly activating Cdx2 (**Fig.S8)**. These results demonstrate NetDes’s ability to optimize small gene circuits that drive cell state transitions, while its performance can depend on quality of the initial GRN.

### Reconstructing sequential cell state transitions in iPSC-to-DE differentiation

So far, we have evaluated the performance of NetDes using synthetic benchmarking tests and simulated biological gene circuit. Next, we will demonstrate its usage by applying NetDes, along with other existing methods, to experimental single-cell RNA-seq data. The goal of this application is to assess the ability of various network inference methods to accurately reconstruct GRNs, while ensuring the inferred networks can capture multi-step cell state transitions observed in the experimental scRNA-seq data.

We analyzed a publicly available time-series scRNA-seq dataset of human induced pluripotent stem cell (iPSC) differentiation toward definitive endoderm (DE)(50). In this experiment, iPSCs from 125 donors (Day 0) were treated with a cytokine-small-molecule cocktail to drive differentiation through mesendoderm (Day 1) to DE (Day 3). Projection of the top 500 highly variable genes (HVGs) onto the first two principal components (PCs) reveals a clear path of continuous cell state transitions from iPSC to DE via mesendoderm (**Fig.3A**). This dataset is ideal for our testing because iPSC differentiation is well studied with extensive literature support, and it allows us to evaluate how well the inferred GRNs capture the multi-step state transitions.

**Fig. 3.**
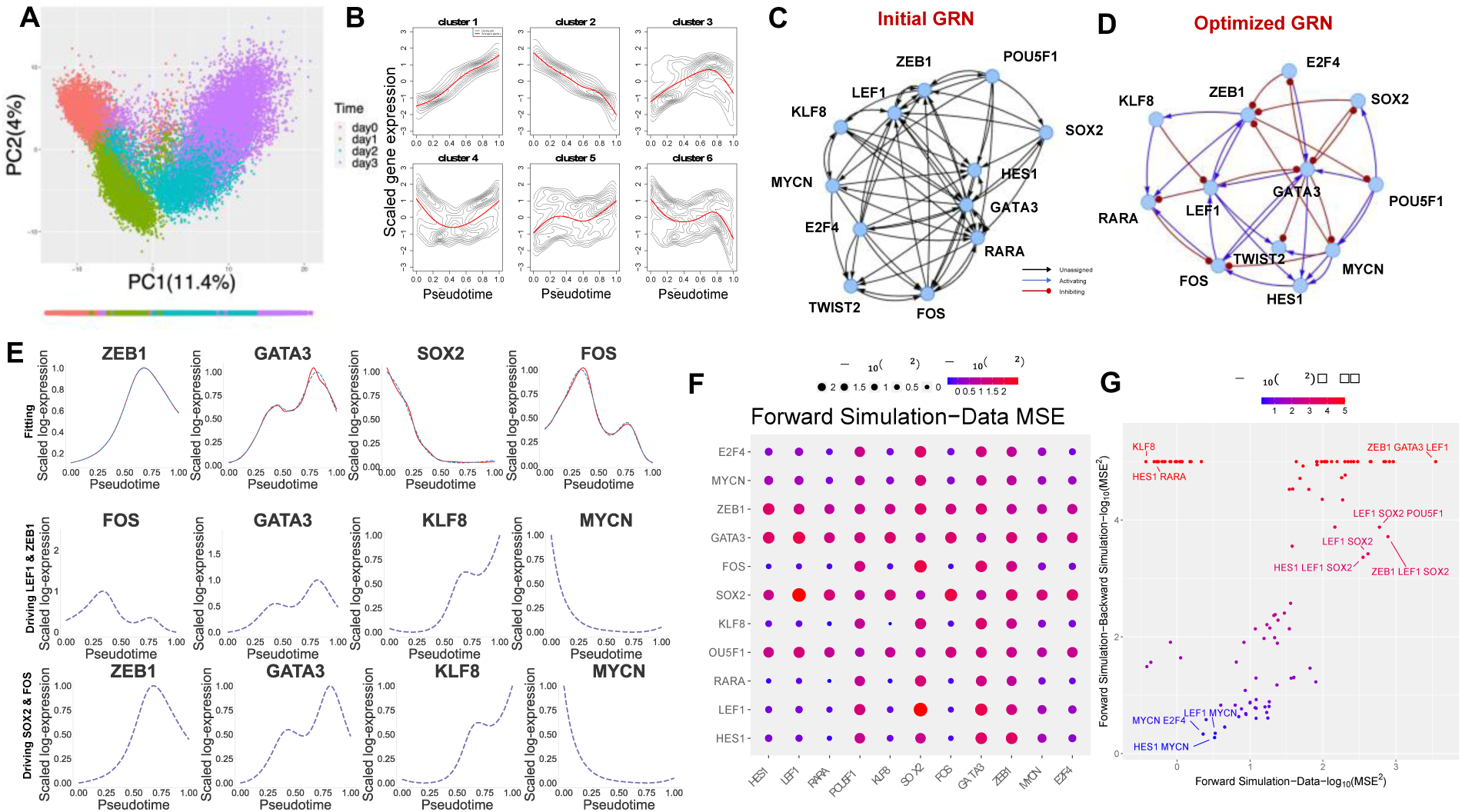
Trajectory inference and GRN optimization for the iPSC-to-definitive-endoderm differentiation. **(A)** Scatter plot showing the normalized expression levels of the top 500 highly variable genes (HVGs) projected onto the first two principal components (PCs). Different colors represent measured time points, as shown for each cell in the scatter plot, with the bottom color bar indicating pseudotime ordering inferred by Slingshot. **(B)** Gene clustering on the top 5,000 HVGs based on the shape of their expression trajectories. Six distinct gene clusters were identified: two with monotonically increasing or decreasing trajectories, two with a single turning point (up-then-down in cluster 3; down-then-up in cluster 4), and two with two turning points. **(C-D)** Network diagrams showing the initial GRN generated using the Rcistarget and TRRUST databases (panel C) and the optimized GRN inferred by NetDes (panel D). In the initial GRN, black lines and arrows represent regulatory edges with unassigned activation or inhibition. In the optimal GRN, the blue lines and arrows indicate activating edges, while the red lines ending in dots represent inhibitory edges. **(E)** The top panel shows comparison of expression trajectories of selected TFs between the transcriptomic data (blue dashed lines) and the simulation from the optimized GRN model (red solid lines). The middle and bottom panels show comparison of simulated gene expression trajectories when the optimized GRN was driven by forward (orange lines) and backward (green lines) signaling on either LEF1 & ZEB1 (middle) or SOX2 and FOS (bottom). In all panels, log-normalized gene expression levels from the data (blue dashed lines) are scaled to a range of 0 to 1, while the simulated expression were scaled accordingly. **(F)** The diagram shows mean squared errors (MSEs) between the experimental and simulated gene expression trajectories, computed for models driven by the expression trajectories of one (diagonal) or two (off-diagonal) TFs. **(G)** Scatter plot summarizing the ability of TFs in driving network gene expression dynamics. The x-axis shows the MSEs between the experimental and forward-simulated gene expression trajectories, while the y-axis shows the MSEs between the forward-simulated and backward-simulated gene expression trajectories. Each dot represents a case when the GRN is driven by one, two, or three TFs.

Before NetDes is applied to build and optimize GRN models, initial data processing and statistical analysis are required to generate an initial GRN. We first inferred pseudotime using Slingshot(56) based on the top 500 HVGs and first five PCs. The inferred pseudotime aligns well with the observed cell state transitions (**Fig.3A**). To mitigate the inherently stochastic nature of gene expression in individual cells and technical noise, such as the dropout effects, we applied PseuodotimeDE(47) to generate smoothed gene expression time trajectories along the inferred pseudotime (details in the **Method** section, examples of smoothed trajectories shown in blue dashed line in **Fig.S3**).

Next, common patterns of gene expression trajectories were identified through trajectory clustering. Conventional clustering methods, such as K-means, K-medoids, and hierarchical clustering, failed to effectively define clusters with distinct trajectory patterns (**Fig.S4**). Thus, we classified the trajectories based on changes in their slope across different time windows (see **Methods**). Applying this approach to smoothed trajectories of the top 5000 HVGs yielded six clusters each showing a distinct gene expression pattern over the pseudotime (contour density maps and the average time trajectories in red lines, as shown in **Fig.3A**). These patterns include trajectories with monotonic increases or decreases, those with a single turning point, and those with two turning points. Trajectories with more than two turning points (15.5%) often correspond to noisy gene expression profiles and were therefore excluded from further analysis.

The characterized time trajectory patterns not only provide insights into the gene expression dynamics during the sequential state transitions in iPSC differentiation but also help to identify potential core regulators of these transitions. Here, we hypothesized that genes in a same trajectory cluster are over-represented by the target genes regulated by a same TF. The core TFs were identified by applying the Fisher’s exact test, where genes within each trajectory cluster were compared with the TF target gene sets from the literature-based NetAct database(57). From this gene enrichment analysis, we identified 16 core TFs enriched in at least one trajectory cluster (q-value < 0.1, as shown in **Fig.S9A**). To reduce false positive TFs resulting from shared target genes, we identified TF pairs with highly overlapping targets (q-value < 10^−20^), and for each pair, we removed the one with the less representative gene expression trajectory (see **Methods** for details). Under this criterion, JUN and TCF7L2 were removed.

The identified TFs include several key stemness drivers, such as SOX2, POU5F1, and MYCN, and TFs involved in epithelial-mesenchymal transition (EMT), including ZEB1, LEF1, KLF8, and TWIST2. The presence of these core TFs suggests an important role of EMT in the differentiation of iPSCs to DE and tight coupling between stemness and EMT regulators. Interestingly, Scheibner et al. also found that SNAI1 couples the loss of stemness with a partial EMT and maintenance of epithelia traits during definitive endoderm formation(75). However, the exact interplays of the stemness and EMT regulators during iPSC-to-DE differentiation are largely undefined. Thus, a systems-biology study on the core GRN of stemness and EMT will provide new insights into the regulatory mechanisms underlying the iPSC-to-DE cell differentiation.

Once the core TFs were identified from the above steps, we connected the TFs by regulatory interactions using both RcisTarget(49) and TRRUST(48) to form an initial GRN (as shown in **Fig.4A**). In the initial GRN, the activating and inhibiting nature of each edge has not been determined yet and will be resolved by model optimization in NetDes. Two TFs ZNF382 and DNMT1 were removed because they are disconnected from the GRN. Note that we have so far illustrated our approach to constructing a reasonable initial GRN of core TFs, while other approaches, such as GENIE3(25), GRNBoost2(76) and DeepSEM(69) could also be applied (with certain adjustment to focus exclusively on core TFs) to accomplish this task.

**Fig. 4:**
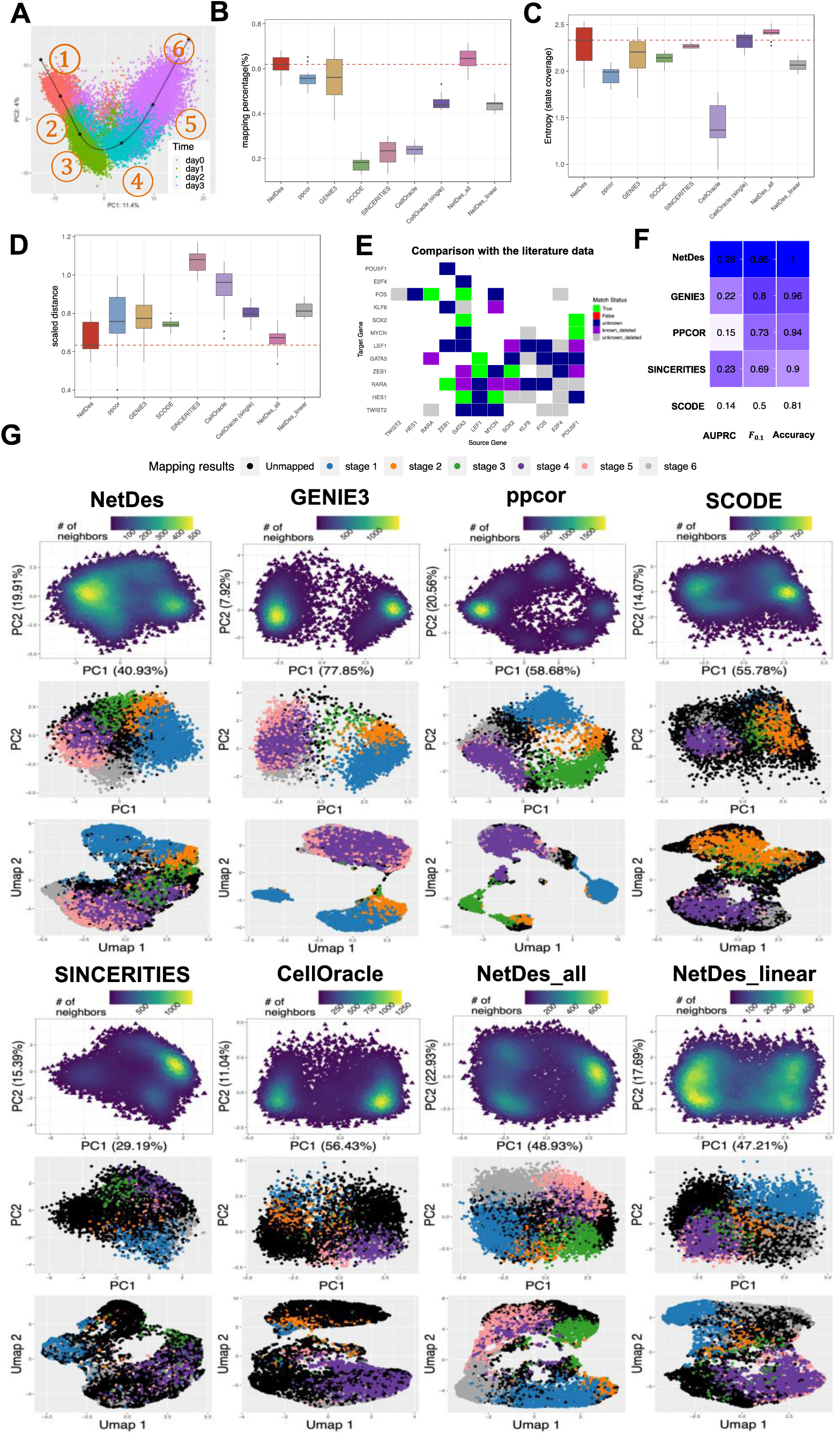
Capability of GRN inference in capturing single cell gene expression states during iPSC-to-DE differentiation. **(A)** Illustration of six reference points along the principal curve computed by Slingshot using the scRNA-seq data projected onto the first two principal components. Panels **B-D** shows the benchmarking of various methods in inferring GRNs that robustly captures observed gene expression states along iPSC-to-DE differentiations. For each inferred GRN containing the specified core TFs and network edges ranging from 31 to 45, a population of single cell gene expression data was simulated by RACIPE and was then compared with the reference gene expression profiles. **(B)** Box plots showing the mapping percentages of simulated gene expression profiles to the reference profiles for each inference method. In this comparison, NetDes_all denotes a variation of NetDes where optimization begins from a fully connected network; NetDes_linear refers to a variation of NetDes which uses a linear ODE model for optimization; CellOracle was applied for both multi-state and single cluster settings (labeled as Cell Oracle single). Tests were performed on 13 GRNs generated by NetDes, 14 by NetDes_linear, and 15 by the other methods. **(C)** Box plots showing the entropy metric that quantifies the disproportion of mapped states. **(D)** Box plots showing the normalized distances in gene expression of unmapped simulated gene expression profiles to the nearest reference profile. **(E)** Comparison of regulatory interactions between the NetDes- inferred GRN (with the best mapping percentage) and the literature data. Each cell in the matrix represents a regulatory interaction from a regulator (source in columns) to a target (target in rows). Different colors represent interactions (1) not present in the initial GRN (blank), (2) predicted and with literature support (green), (3) with literature support but predicted with wrong activation/inhibition nature (red), (4) predicted but without literature evidence (dark blue), (5) with literature support but removed by NetDes (purple), (6) not predicted and no literature evidence (gray). **(F)** Performance of GRN inference methods in predicting literature-based regulatory interactions. Interactions inferred by a method was filtered to retain only interactions from the initial GRN before performance evaluation. The heatmap and numbers indicate the AUPRC, accuracy, and F_0.1_. **(G)** Simulated gene expression profiles for the inferred GRNs with the highest mapping percentage for each method. Each set of subplots shows the density plot of simulator gene expression projected onto the first two principal components (top), the scatter plot of gene expression colored by their mapped reference states on the PCA space (middle), and the same scatter plot but projected onto the first two UMAP dimensions (bottom).

### NetDes builds a dynamical model for the GRN driving iPSC-to-DE differentiation

Next, we applied NetDes to optimize both the GRN structure and associated dynamical model, using the initial GRN (12 core TFs and 59 candidate interactions) and gene expression time trajectories. For each target gene (also a TF), we optimized an ODE model by using the gene expression trajectories of the target and its candidate regulators as specified in the initial GRN (**Fig.3C**). After optimizing models for each core TF, we assembled regulatory interactions into an integrated GRN. Depending on the choice of the optimization threshold cutoffs, we built GRNs of different sizes, ranging from 31 to 45 interactions. The best performing GRN contains 36 interactions, as shown in **Fig.3D** (see the next section for details on its selection). For most TFs, this approach successfully optimized the ODE models that generate comparable trajectories to those from the observed data (top panel in **Fig.3E** and **Fig.S10**). A few TFs, such as KLF8 and E2F4, showed mismatches between fitted and observed trajectories, possibly due to the complexity of their expression dynamics or missing upstream inputs. Overall, the ODE-based optimization accurately reproduces individual genes’ expression even after removing 23 candidate interactions from the initial GRN.

After optimizing the ODE for each TF, we assembled the full set of ODEs for the entire core TF GRN to evaluate whether the gene expression dynamics of the whole system could still be captured by the model. This evaluation is nontrivial because complex regulatory interactions, such as coupled feedback and feedforward motifs, can substantially influence the overall dynamics. Moreover, because iPSC-to-DE differentiation was experimentally induced by a cocktail of factors(50) that directly regulate TFs, such as EGR1 and members of the SOX and SMAD families, we simulated the GRN dynamics by driving the system with one, two, or three TFs using their expression trajectories to mimic extrinsic signaling inputs. These simulations allow us to evaluate the extent to which TF expression trajectories within the GRN can be driven by extrinsic signaling and to identify which TFs are likely to drive iPSC-to-DE differentiation.

We then asked whether specific TF combinations could externally drive the optimized GRN through the differentiation trajectory, thereby identifying candidate drivers of the transition. We setup a series of simulations by considering all possible combinations of one or two TFs as drivers. For each combination, we simulated the expression trajectories of the remaining TFs by driving the selected TFs according to their trajectories along the pseudotime to induce cell differentiation (referred to as forward signaling). Additionally, we simulated the reverse process by driving the selected TFs with their trajectories along the reverse pseudotime (referred to as backward signaling). Middle and bottom panels in **Fig.3E** and **Figs.S11, S12** show the simulated gene expression trajectories for both forward (orange lines) and backward (green lines) signaling, compared to the observed trajectories from the data (blue dotted lines), for two representative signaling combinations: one driven by LEF1 and ZEB1, and the other by SOX2 and FOS. For LEF1 and ZEB1 (middle row in **Fig.3E** and **Fig.S11**), several TFs displayed distinct trajectories between forward and backward signaling, indicating the presence of hysteresis, a characteristic feature of multistable dynamical systems. As a result, the simulated gene expression trajectories tended to deviate from the observed data, suggesting that the GRN may encounter barriers in the regulatory landscape when driven by these TFs. We also found that in some TFs, the simulated trajectories exhibited reduced fold changes in gene expression compared to the data, likely due to attenuation through multiple regulatory steps encoded in the coupled ODEs; however, the overall shape of the trajectories was often preserved after appropriate scaling of the observed data. In contrast, for the combination of SOX2 and FOS (bottom panel in **Fig.3E** and **Fig.S12**), the simulated trajectories from forward and backward signaling closely matched, indicating the absence of hysteresis. In this specific case (although not necessarily for all non-hysteretic combinations), the simulated trajectories aligned more closely with the observed data. This suggests that driving the GRN with SOX2 and FOS may create a smoother path in the regulatory landscape for inducing cell state transitions.

To systematically evaluate all combinations of one or two TFs, we plotted heatmaps of the mean square errors (MSEs) between forward-simulation trajectories and the data (**Fig.3F**), as well as the MSDs between forward and backward simulations (**Fig.S9B**). **Fig.3G** summarizes both metrics in a scatter plot with annotations on representative two or three TF combinations. Simulations revealed that single-TF drivers often led to hysteresis and were generally insufficient to fully recapitulate the observed gene expression dynamics, though stemness-associated TFs like SOX2 and MYCN performed better than others. In contrast, dual-TF combinations, particularly those involving ZEB1, SOX2, or GATA3, more effectively drove the GRN without hysteresis, suggesting a smoother landscape when driving the transitions. Remarkably, GATA3 is a known driver of the embryonic stem cell differentiation(77). When used to drive the GRN alone, GATA3 generated favorable results. Furthermore, combining GATA3 with EMT genes such as LEF1, KLF8, and ZEB1 improved the driving performance (e.g., **Fig.S13**). In addition, certain TFs in the system, such as FOS, TWIST2, RARA and E2F4, cannot effectively drive the cell state transitions. Together, the driving simulations show that the optimized GRN can be externally driven to capture the desired cell state transitions, while also identifying candidate drivers for the system.

### NetDes performs competitively in iPSC-to-DE GRN inference

Next, we evaluated the performances of several GRN inference methods in reconstructing regulatory networks capable of recapitulating gene expression states during iPSC-to-DE differentiation, using both GRN simulation and literature-based evidence. NetDes was benchmarked against five other methods: GENIE3(25), ppcor(22), SCODE(30), SINCERITIES(23), and CellOracle(31). For each method, GRNs of the identified core TFs were inferred with varying number of regulatory interactions, ranging from 31 to 45. This selection ensures the constructed GRNs are likely to be connected and not too dense. We also test two variations of NetDes: one using linear ODE models for optimization (denoted as *NetDes_linear*) and another in which optimization started from a fully connected TF network (denoted as *NetDes_all*). For each inferred GRN, we applied RACIPE (using sRACIPE(67) package), which generates gene expression profiles by simulating ensembles of ODE models based on the same network topology but with randomly sampled kinetic parameters. Previous studies have shown that RACIPE captures network states that are robust to parameter variations(65–68), which may mimic cell-to-cell variability and gene expression noise, and that simulated gene expression profiles can be effectively compared with experimental scRNA-seq(78) . Moreover, mathematical modeling using RACIPE was applied to each GRN to ensure that performance evaluation does not depend on the optimized ODE models in NetDes (with fixed kinetic parameters) and treats all methods equally by using randomized parameters.

In the case of iPSC-to-DE differentiation, we mapped simulated gene expression profiles from RACIPE to six representative gene expression snapshots (denoted as the reference cell states), evenly selected along the principal curve of the scRNA-seq data (**Fig.4A**), to evaluate how well each GRN topology captures cell states along the continuous differentiation trajectory. Also, we calculated the entropy of the mapped state distribution mappings to assess how evenly each GRN captures the continuous cell state transitions (see **Methods** for details). NetDes achieved the highest weighted mapping percentage and entropy among all tested methods, regardless of whether the optimization starts from a literature-based or fully connected initial GRN (**Figs.4BC, Fig.S14**). The mapping percentages were drastically reduced when linear models were used in NetDes, suggesting the need of nonlinear ODEs for better network modeling. Moreover, for simulated gene expression profiles that could not be mapped to any reference cell state, those derived from NetDes-inferred GRNs were overall closer to these reference cell states than those from other methods (**Fig.4D**). In contrast, GRNs inferred by other methods often produced clusters of simulated gene expression profiles that significantly deviated from the observed data. These results demonstrate that GRNs inferred by NetDes more effectively capture different cell states during differentiation.

For each method, we chose the GRN with the highest mapping percentages and visualized the densities and mappings of RACIPE-simulated gene expression profiles using PCA and UMAP projections (**Fig.4G**). Simulated profiles from NetDes-inferred GRNs tended to form more continuous trajectories across different reference states in the low dimensional space. In contrast, GRNs inferred by other methods often produced simulated profiles that formed separated clusters, or exhibited incorrect ordering of mapped states.

To further evaluate the influence of the initial GRN in the benchmarking, we filtered the inferred regulatory interactions for each method based on the same initial GRN. The only exception was CellOracle, which incorporates prior information from its own prebuilt base GRN rather than the initial GRN. With this adjustment, we observed performance improvements in mapping percentages in certain methods (**Fig.S15**). NetDes still outperformed all other methods in the benchmark. Overall, RACIPE simulations and state mapping analyses show that NetDes consistently outperforms other methods, regardless of whether it starts from a literature-based initial GRN. Nonetheless, we recommend NetDes optimization starting from an initial GRN containing substantially fewer edges, to make model inference more realistic and to reduce computational cost.

Lastly, we compared the TF GRNs inferred by various methods against literature evidence (**Table.S1**). We found NetDes outperformed all other methods when evaluating the AUPRC, accuracy, and *F*_0.1_ scores (**Figs.4EF** and **Figs.S16, S17**). Together, these results indicate NetDes as the top-performer in constructing robust and accurate GRN models capturing multi-step cell state transitions.

### GRN coarse-graining reveals a stemness-intermediate-EMT circuit logic

So far, we have applied NetDes to construct a core TF GRN driving iPSC-to-DE differentiation (**Fig.3D**, with the highest mapping percentage). The GRN reveals detailed TF interactions, but how they drive differentiation together remains vague due to the complexity of the GRN. We applied SacoGraci(44) to coarse-grain the GRN into a small circuit motif where nodes represent TF groups and edges represent the collective effects of gene regulation among gene groups.

According to 10000 simulated gene expression profiles of the full GRN using RACIPE (leftmost panel in **Fig.5B**), we clustered TFs into three groups: (1) SOX2, POU5F1, MYCN, TWIST2, and E2F4, as stemness genes with early expression; (2) FOS, LEF1, and HES1 and GATA3, as genes expressed during the intermediate states; (3) ZEB1, KLF8 and RARA, as genes associated with EMT with late expression. Note that TWIST2, although known to be an EMT regulator, was found to have decreasing expression during the differentiation process, and thus was clustered in Group 1. Also, we set the cluster of models into three based on the RACIPE simulated profiles. With the gene and model grouping, we performed 30 iterations of SacoGraci to construct the best coarse-grained (CG) circuit (right panel of **Fig.5A**) according to the consistency of simulated gene expression profiles of the CG circuit and the full GRN. We found the network interactions from the CG circuit and the full GRN were highly consistent. Furthermore, we found the simulated gene expression distributions for the full GRN, those of the CG circuit, and the profiles from the experimental scRNA-seq data (heatmaps of simulated profiles shown in **Fig.5B**, and density maps shown in **Fig.5C**) are qualitatively similar, suggesting a successful coarse-graining of the CG circuit. Interestingly, the optimal CG circuit reveals mutual inhibition between the stemness TFs (Group 1) and EMT TFs (Group 3). In addition, intermediate TFs (Group 2) form negative feedback with the stemness TFs, but positive feedback with the EMT TFs. During iPSC-to-DE differentiation, stemness TFs (Group 1) first activate intermediate TFs (Group 2). As Group 2 expression increases, these intermediate TFs in turn induce EMT TFs (Group 3). Groups 2 and 3 then engage in mutual positive feedback while gradually repressing stemness TFs (Group 1). Such repression in turn reduces the activities of the Group 2 genes in the later stage through the positive regulation from stemness genes. In summary, GRN coarse-graining uncovers the potential regulatory logic governing multi-step iPSC-to-DE differentiation.

**Fig. 5:**
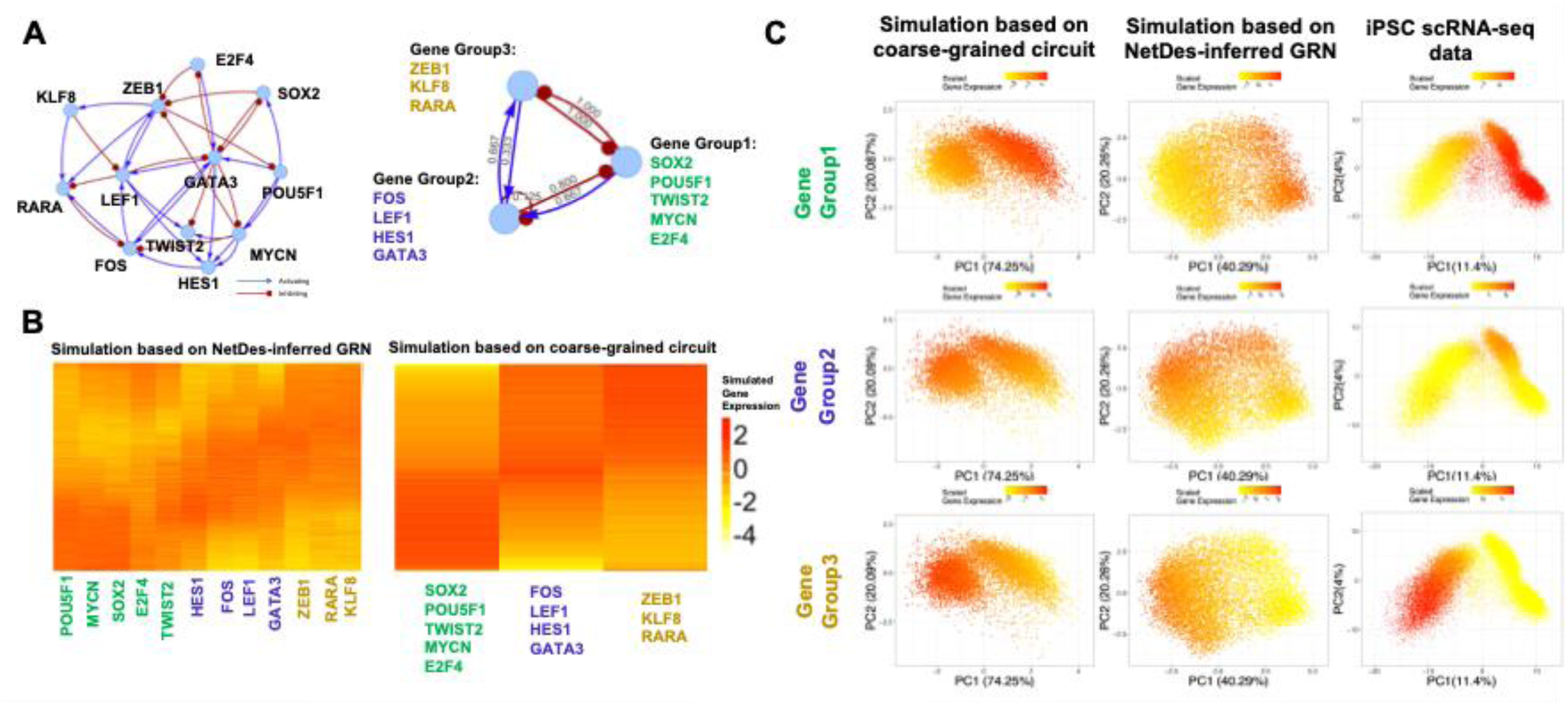
GRN coarse-graining to identify regulatory motif. **(A)** Illustration of the NetDes-inferred full GRN and the corresponding coarse-grained (CG) circuit diagrams inferred by SacoGraci. In the CG circuit, the TFs from the full GRN are grouped into three CG nodes based on expression similarity: Group 1 (SOX2, POU5F1, TWIST2, MYCN, E2F4), Group 2 (FOS, LEF1, HES1, GATA3), and Group 3 (ZEB1, KLF8, RARA). Numbers along each regulatory edge in the CG circuit represents the ratio of activating/inhibitory regulatory interactions present in the full GRN. **(B)** Heatmaps of simulated gene expression for the NetDes-inferred GRN (left) and CG circuit (right). Rows are ordered by inferred pseudotime from Slingshot using the first two principal components of the simulated gene expression profiles. **(C)** Comparison of average gene expression levels for each CG group for the simulated gene expressions from the CG circuit (leftmost column), the simulated gene expression from the full GRN (middle column), and the experimental scRNA-seq data (rightmost column). Each row shows scatter plots of gene expression projected onto its first two principal components, where colors represent mean expression levels for genes of each group. Columns represent simulations from the CG circuit (leftmost), simulations from the TF GRN (middle), and experimental iPSC-to-DE differentiation scRNA-seq data using the top 500 highly variable genes (rightmost).

### NetDes models diverse continuous cell-state-transition systems

To showcase NetDes’ usage in modeling diverse continuous cell-state transitions systems, we applied the similar workflow to three additional time-series scRNA-seq datasets representing distinct biological process: epithelial-mesenchymal transition (EMT) in A549 cells(51), erythroid differentiation (ERY)(52) and dendritic-cell differentiation (DC)(53). For each system, we inferred pseudotime, identified core TFs, constructed an initial GRN, and then applied NetDes to infer GRN and corresponding ODE models following the same procedure used for the iPSC-to-DE differentiation data.

Importantly, the three datasets differ in the density of their initial GRNs (i.e., the average number of edges per TF), allowing us to assess the performance of NetDes across initial networks of varying connectivity. The initial GRN for EMT is a fully connected network (without self-regulation) of eight core TFs. In contrast, the initial GRNs for ERY and DC are sparser: 15 TFs with 100 edges (about half of all possible TF–TF edges) for ERY, and 19 TFs with 72 edges (about one-fifth of all possible edges) for DC. Despite this wide range of initial-GRN densities, NetDes robustly inferred dynamical GRN models in all three systems, indicating that its performance is robust to variation in the size and edge density of the initial network.

Across all three systems, the cells formed continuous gene expression trajectories in principal component space consistent with their measured time points (**Fig.6A**), and NetDes inferred a GRN of core TFs, along with a dynamic model, for each system (**Fig.6B)**. Using RACIPE to simulate gene expression states from each optimized GRN, we found that the simulated states successfully captured the observed cell-state distributions, with the experimental expression trajectory, projected onto the same principal components, connecting the starting and ending states along the biologically expected direction of the transition (**Fig.6C**). For the GRN, its ODE model reproduced the expression dynamics both in fitting and gene-driving simulations, quantified by the Nash– Sutcliffe efficiency (NSE) between simulated and reference trajectories, which we found good agreements (the NSE exceeding 0.5) for 39 of 42 genes in the fitting simulations and 41 of 42 genes in the gene-driving simulations across the three systems (**Fig.S18**). We further examined the capacity of the inferred networks to drive the observed transitions by calculating the MSE between the experimental and forward-simulation trajectories and between the forward and backward simulation trajectories, identifying TF combinations that most effectively drive each transition (**Fig.6D**). Remarkably, the top driving TF combinations recover known regulators of each transition. For the EMT, the top drivers are HMGA1 together with ETS1 and KLF2, consistent with HMGA1 and ETS1 as promoters of TGF-β–induced EMT (79,80). For the erythropoiesis, the top drivers are GATA1 pairs, such as GATA1–RARA and GATA1–XBP1, reflecting the role of GATA1 as the master erythroid regulator and its antagonism with FLI1(81,82). For the dendritic cell differentiation, we identified RELB, RARA and FLI1, consistent with the requirement of RELB for cDC2 development and retinoic-acid control of cDC2 fate(83,84) (**Fig.6D**). Notably, in all these cases, the top driver predictions recover known master regulators without prior knowledge. Finally, we benchmarked NetDes against other GRN-inference methods using the same metrics as we used for the iPSC data. Across all three datasets, no other method matched NetDes on all three metrics at once: a higher median mapping percentage, a higher median entropy of mapped states, and a lower normalized gene-expression distance for unmapped states (**Fig.6E**). In the case of DC differentiation, GENIE3 achieved the largest mapping percentage but the lowest entropy of mapped states, suggesting its GRN ensemble collapsed onto a single cell state; SCODE, by contrast, had the highest entropy but a low mapping percentage. Overall, those results show that NetDes robustly reconstructs GRNs with dynamic models across different continuous cell-state-transition systems.

**Fig. 6.**
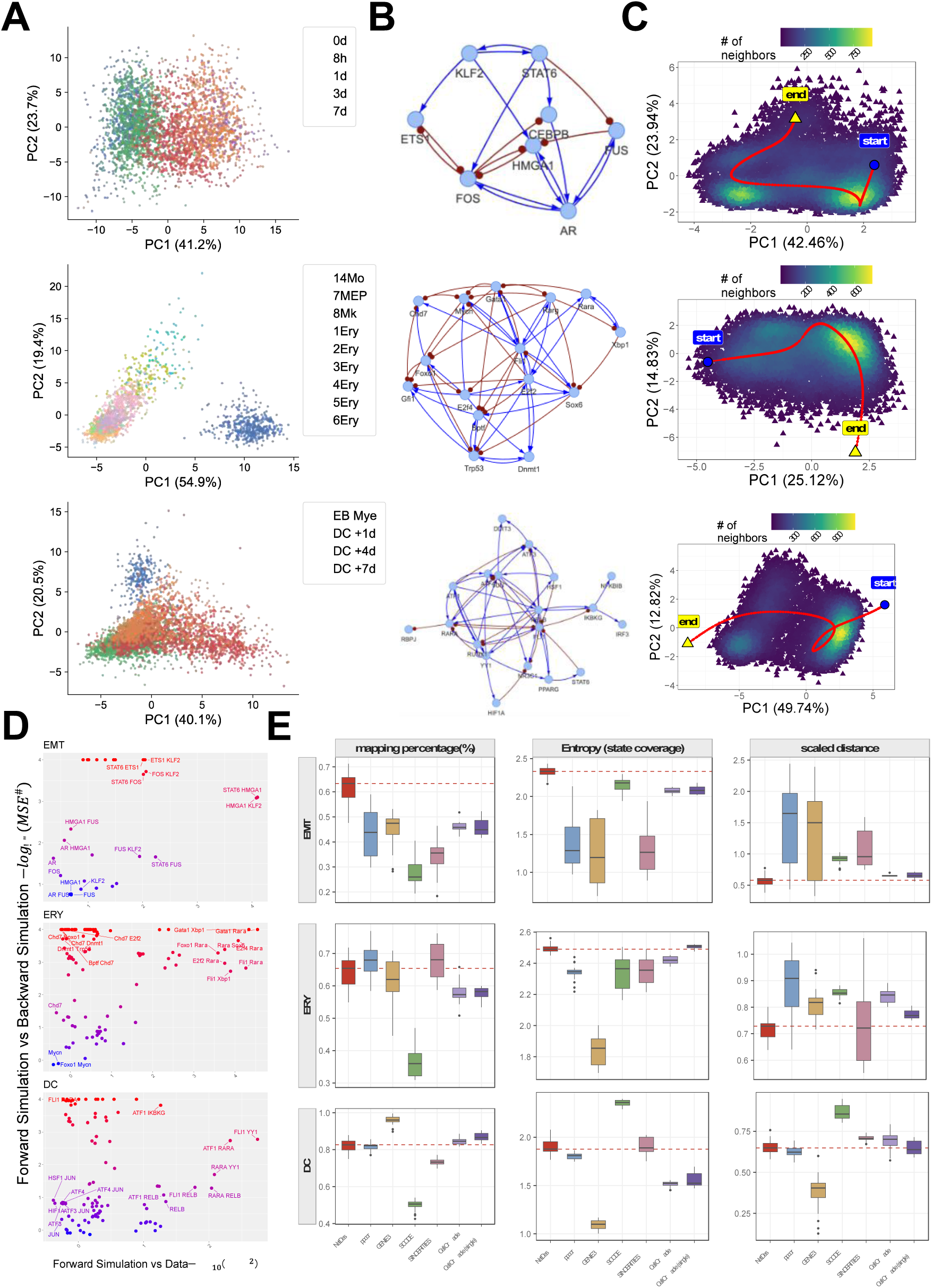
NetDes reconstructs GRN with dynamic models across another three cell-state-transition systems. NetDes was applied to three different time-series scRNA-seq datasets: epithelial-mesenchymal transition (EMT), erythropoiesis (ERY), and dendritic-cell differentiation (DC), shown in the top, middle and bottom rows in each panel, respectively. **(A)** Scatter plots of cells projected onto the first two principal components for each dataset, colored by the measured time points. **(B)** NetDes-inferred GRN of the core TFs for each dataset. Blue arrows indicate activating edges and red lines ending in dots indicate inhibitory edges. **(C)** Density plots of RACIPE-simulated gene expression projected onto the first two principal components, the red curve shows the smoothed expression trajectory of the core TFs, inferred by PseudotimeDE from the scRNA-seq data and projected onto the same PCs, connecting the starting (start) and ending (end) cell states. **(D)** Scatter plot summarizing the ability of TFs to drive network gene expression dynamics for each dataset. The x-axis shows the MSE between the experimental and forward-simulated trajectories; the y-axis shows the MSE between the forward- and backward-simulated trajectories. Each dot represents a driving case in which the GRN is driven by one, two, or three TFs, colored by MSE. **(E)** Comparison of NetDes against other GRN-inference methods (ppcor, GENIE3, SCODE, SINCERITIES and CellOracle) across the three datasets. CellOracle was applied for both multi-state and single cluster settings (labeled as Cell Oracle single). For each dataset (in rows), box plots show the weighted mapping percentage (left columns), the entropy of the mapped state distribution (middle columns), and scaled distance (right columns) of the RACIPE-simulated states to the reference profiles. NetDes is highlighted in red and the dashed line marks the NetDes median in each panel. A higher mapping percentage, higher entropy, and a lower scaled distance indicate a better agreement between the simulated ensemble and the observed cell states from the experimental data.

## Discussion

In this study, we developed a computational method, NetDes, that infers core transcription factor (TF) regulatory networks driving continuous, multi-step cell state transitions using single cell gene expression trajectories. NetDes integrates top-down and bottom-up systems biology by fitting ordinary differential equation (ODE) models to observed gene expression trajectories. The resulting dynamic models are interpretable and allow further simulation and analysis of regulatory systems. We evaluated the performance of NetDes through extensive benchmarking on synthetic gene expression trajectories, a simulated biological gene circuit, experimental single-cell RNA-seq data from the BEELINE collection, and time-series scRNA-seq datasets from four different kinds of cell-state-transition systems (iPSC-to-DE differentiation, EMT, erythropoiesis, and dendritic-cell differentiation).

A major advantage of NetDes is its ability to identify a minimal gene regulatory network (GRN) of core TFs that drives observed cell state transitions. By explicitly modeling gene expression dynamics using ODEs, NetDes goes beyond static or association-based network inference and enables systems-biology simulation of GRN dynamics under network perturbations or extrinsic signals. These in silico experiments and explorations facilitate the generation of new testable hypotheses for experimental validation, which have the potential to improve causal inference and predictive modeling of regulatory systems. Moreover, simulations for the GRNs driven by combinations of interactions rather than individual edges may help find some synergistic effects overlooked by traditional methods. Compared to existing methods, NetDes utilized a systems-biology dynamical modeling approach that explicitly constructs ODE models to capture the gene expression dynamics during observed multi-step cell state transitions. Moreover, in benchmarking, we evaluate how well a GRN covers cell state populations using ensemble-based simulations. Such a test is independent of model fitting or statistical inference of GRNs, and could serve as a general approach for model evaluation or selection.

Despite its advantages, NetDes has several limitations. First, the current method requires high-quality pseudotime or trajectory inference, and its accuracy can affect the performance of network modeling. Second, the model optimization assumes that a deterministic ODE model is sufficient to capture gene expression dynamics, which may perform less well in cases where stochastic dynamics or cell-cell communication are significant. In such scenarios, ensemble-based modeling approaches, such as RACIPE, may better capture these effects. Third, the network optimization step in NetDes is less efficient when a TF has more than ten candidate regulators. This issue can be alleviated through parallelization and the integration of compiled code for optimization. Fourth, the current implementation of NetDes assumes a single, continuous trajectory and does not accommodate branching or bifurcating trajectories associated with multiple cell fates. We plan to extend the method to such scenarios in the future. Fifth, synthetic trajectory generation, model fitting, and ensemble-based GRN simulations all rely on Hill-kinetics ODE models. While a such modeling choice is widely used in the systems-biology literature for gene circuit and network modeling(40,85,86), we have not yet evaluated how alternative mechanistic formulations might affect method performance, which is worth further investigation. Finally, because all genes were assigned a common degradation rate to maintain quasi-steady-state dynamics, the model captures trajectory shape and driver-gene logic but not the true timescale of the transition. Such a detailed model would require specification of gene-specific half-lives, which remains an open challenge to be addressed. Beyond addressing these limitations, future work could extend NetDes to model different types of gene regulation, such as posttranscriptional regulation, and to generalize the method for multi-omics data, such as transcriptomic, epigenomic, and proteomic measurements, to enable more comprehensive modeling.

In conclusion, NetDes provides a general approach for mechanistic modeling of transcriptional regulatory networks driving cell state transitions. By combining core TF network inference with ODE-based model optimization and simulations, it offers new opportunities to elucidate design principles of cell fate regulation and to develop effective strategies for driving cell state transitions.

## Code availability

NetDes package is available at both GitHub https://github.com/lusystemsbio/NetDes and PyPI https://pypi.org/project/NetDes/. Analysis scripts, including in-silico benchmarking, scRNA-seq application and TF regulatory network modeling, are available at https://github.com/lusystemsbio/NetDesAnalysis/.

## Data availability

The data used in this study are as follows. The iPSC scRNA-seq data are available at https://zenodo.org/records/3625024#.Xil-0y2cZ0s. The EMT scRNA-seq data are available from the Gene Expression Omnibus under accession GSE147405. The erythropoiesis scRNA-seq data are available from the Gene Expression Omnibus under accession GSE72857. The human in vitro myelopoiesis (DC) scRNA-seq data are available from ArrayExpress under accession E-MTAB-11623. The hESC and hHep scRNA-seq datasets used for benchmarking, together with the non-specific, hESC and HepG2 ChIP-seq networks used to define the initial GRNs and the ground truth, are from the BEELINE collection and are available at https://zenodo.org/records/19009603. The orginal scRNA-seq data are available from the Gene Expression Omnibus under accessions GSE75748 (hESC) and GSE81252 (hHep). The synthetic benchmark datasets are available at https://github.com/lusystemsbio/NetDesAnalysis/. The RcisTarget human motif database is available at https://resources.aertslab.org/cistarget/databases/homo_sapiens/hg19/refseq_r45/mc9nr/gene_based/. The literature-based human TF–target database from NetAct is available at https://github.com/lusystemsbio/NetAct/blob/master/data/hDB.rdata.

## Acknowledgements

1. Y. You, C. Caranica and M. Lu are supported by startup funds from Northeastern University, by the National Institute of General Medical Sciences of the National Institutes of Health under Award Number R35GM128717. Y. You and M. Lu acknowledge their affiliation with the Center for Theoretical Biological Physics at Northeastern University and appreciate the support provided by the center. The authors used Claude to check grammar and improve clarity of the manuscript text. The authors reviewed and edited all AI-assisted output and take full responsibility for the content of the manuscript.

