## Supplementary Information (Methods, Figures, and Tables) for "Building dynamical models of multi-step state transitions from single cell gene expression trajectories"

#### Supplementary Materials

#### Supplementary Methods

##### Detailed four-stages of model optimization

*Initial model optimization (first stage)*

To ensure robust model fitting and prevent the nonlinear fitting from being trapped in local minima, we devised an approach consisting of four stages for model optimization (**Fig.S1A**). During the first stage, we performed an initial round of optimization without regulation. First, a substantial number of initial parameters were sampled for the optimization as follows. For each gene $i$ and its regulator j, we sampled the initial $\lambda_{ij}$ from $4 > \lambda_{ij} > 2$ (for activation) and $0.5 > \lambda_{ij} > 0.25$ (for inhibition) to ensure all activation/inhibition combinations from all regulators of gene $i$. In addition, we uniformly sampled the initial $R_{ij}$ from $(\, 0.3 \cdot \max(y_j), 0.7 \cdot \max(y_j))$, the initial $n_{ij}$ from $(0.01, 0.99)$, and the initial $C$ from (-2,2). This approach results in a total of $2^{N_i}$ sets of initial ODE parameters, where $N_i$ is the number of regulators of gene $i$. Second, starting from each of the $2^{N_i}$ sets of initial parameters, we applied five-fold cross-validation (CV) to fit the model using **Eq. (3)** but without the regulation terms ($w_i = 0$) (**Fig.S1A**, step 1). For the five-fold CV, we separated the data evenly into five parts based on the pseudotime and, for each iteration, used four parts to fit the model and the rest one as test data to calculate the mean squared errors (MSEs). After the five-fold CV, we computed the mean MSE as the final MSE under this parameter set. Third, among all the fitted models, we selected the models with the best 50% (if $N_i \leq 6$) or the top 50 (if $N_i > 6$) models in MSEs, and then we re-optimized these models starting from resampled initial parameters (but keeping the same $\lambda_{ij}$ ranges) again for ten times to further improve the fitting outcomes (**Fig.S1A**, step 2). Finally, the top 100 models in MSEs (or all top 50% models if the total number is less than 200) will be used later for a second stage of model optimization where regulation terms were added (described in the next section).

*Tuning parameter estimation and optimization with regulation (second stage)*

Right after the first stage of model optimization without using the regulation terms, we estimated the tuning parameters so that the regulation terms on the maximum fold change parameters could be properly incorporated during the second stage (**Fig.S1A**, step 3). For each gene $i$ , we selected the initial parameters from the top four fitted models according to the MSE values (see the 1st stage). We selected a series of tuning parameters $w_i$ (a total of 20 numbers by default) by evenly sampling $\ln(w_i)$ from -15 to 2. For each tuning parameter, we performed more model fitting using **Eq. (3)** and with the regulation term by a five-fold CV test. From the plot of $w_i$ (x-axis) *v.s.* $\log(mean(MSE))$ (y-axis, the average is over the above-mentioned four fitting cases), we fitted a spline curve and determined the tuning parameter $w_i$ by the minimum of the spline curve (**Fig.S2D**). This procedure was repeated to determine the tuning parameter for every gene.

From the outcomes of the 1st stage model optimization, we then performed a second stage optimization by incorporating the regulation terms as follows (**Fig.S1A**, step 4). The top 100 models in MSEs (or top 50% models if the total number of models is less than 200) were optimized again with the regulation terms. Here, for each model, its initial parameters were re-sampled while keeping the same patterns of activating/inhibiting edges (*i.e.*, $\lambda_{ij}$ were re-sampled according to the same ranges as specified in the 1st stage). After this round of optimization, the optimized models were ranked again according to the MSEs. Incorporating regulation for the fold change parameters alleviates potential overfitting by effectively reducing the number of regulators for each gene.

*Optimization on reduced regulatory scenarios (third stage)*

To further improve the robustness of optimization and determine a minimal regulatory model recapitulating gene expression trajectory, we employed a third stage optimization on models where subsets of regulators were selected for additional optimization but without regulation. First, from the optimized models in the previous step (step 4), we identified all *removable* incoming edges for each gene $i$ (**Fig.S1A**, step 5). Here, we considered an edge to be removable, if the optimized $\lambda_{ij}$ can be both $\lambda_{ij} > 1$ and $\lambda_{ij} < 1$ within all the optimized models by step 4. Second, we generated all possible regulatory scenarios for gene $i$, where a subset of removable incoming edges were deleted. For each of such scenarios, we fitted the ODE model again ten times, where the new optimization starts by using the fitted parameters from the top 10 (instead of top 100 to reduce computational cost) optimized models from step 4. In this fitting step, we did not use regulation terms and cross validation. Newly fitted models and their corresponding regulatory scenarios were ranked by a reducibility index $I$, defined by

$$I = \sum_{h=1}^{10} \frac{MSE_h}{MSE_{ori,h}} \quad \text{(S1)},$$

, where $MSE_h$ are the optimized MSE under different initial parameters, and $MSE_{ori,h}$ is the original MSE from the $h^{th}$ model in the top 10 model optimization in step 4.

The reducibility index highlights the impact of edge deletions on model fitting. For the regulatory scenarios whose reducibility indices are below a specific cutoff value, we determined the optimal regulatory scenario according to the fewest number of edges. If there are multiple of such scenarios, the optimal scenario was selected to be the one with the lowest reducibility index. By selecting a specific cutoff value, we obtained optimal regulatory scenarios for each gene and built a GRN by assembling regulatory interactions from all genes. By choosing different cutoff values, we can generate GRNs of varying sizes.

*Optimization to refine and finalize the ODE model (final stage)*

Once we constructed a GRN corresponding to a specified reduced regulatory scenario, we finalized a corresponding ODE model that specifies the activation/inhibition nature of each interaction using the final optimization steps (**Fig.S1B**, step 8). First, an initial optimization was performed without regulation for all optimized models in step 4 but under the reduced regulatory scenario (**Fig.S1B** step 9, like **Fig.S1A** step 1). Here, the initial parameters of the reduced models were derived from the corresponding fitted parameters of these full models. Second, the top four fitted models from the previous step were used to estimate the tuning parameters (**Fig.S1B** step 10, similar to **Fig.S1A** step 3). Third, another round of optimization was then performed for all models in step 9 but with regulation terms (**Fig.S1B** step 11, similar to **Fig.S1A** step 4). Finally, we identified a subset of the optimized models with the most frequent activation/inhibition pattern of the regulators according to the signs of $\lambda_{ij}$. From these models, we identified the best model according to the smallest MSE (**Fig.S1B** step 12). This step generated the final ODE model of gene $i$ . The process was repeated for each gene to complete the dynamical model of the whole GRN.

#### Simulation of a gene circuit driving ICM-to-TE transition

To simulate the gene expression dynamics of the gene circuit driving ICM-to-TE transition, we set the expression trajectory of the signaling gene Fgf ($x_4$) as an input signal changing from its initial value of 25 (arbitrary unit) to its final value of 150 over the course of the simulation.

$$x_4(t) = \begin{cases} 25 + 125 * \dfrac{1 - \cos\left(\dfrac{\pi\, t}{T/2}\right)}{2}, & 0 \le t \le T/2 \\ 150, & T/2 < t \le T \end{cases} \quad (S2),$$

where T is the total simulation time, which is set to 1000.

For other three genes Oct4 ($x_1$), Cdx2 ($x_2$) and Esrrb ($x_3$), we simulated their gene expression dynamics according to:

$$\frac{dx_i}{dt} = g_i \prod_{j \neq i} \left\{ \lambda_{ij} + \frac{(1 - \lambda_{ij})}{\left[1 + \left(\frac{x_j}{R_{ij}}\right)^{n_{ij}}\right]} \right\} - k_i x_i \quad (S3),$$

The parameter set for the simulation is shown in **Supplementary Table 2**.

#### Synthetic dataset for benchmarking

To evaluate the performance of network inference, we generated synthetic datasets of time-series gene expressions for regulators and target genes. We randomly generated ten curves from polynomials of either fourth or fifth degrees to represent the gene expression trajectories of ten genes (**Eq. S4**).

$$\begin{aligned}
I_1(x) &= \frac{1000 + x \cdot (x - a_0) \cdot (x - a_1) \cdot (x - a_2) \cdot (x - a_3)}{100} \\
I_2(x) &= \frac{1400 - (x - a_4) \cdot (x - a_5) \cdot (x - a_6) \cdot (x - a_7) \cdot (x - a_8)}{100} \\
I_3(x) &= \frac{5500 + (x - a_9) \cdot (x - a_{10}) \cdot (x - a_{11}) \cdot (x - a_{12}) \cdot (x - a_{13})}{100} \\
I_4(x) &= \frac{1000 + (x - a_{14}) \cdot (x - a_{15}) \cdot (x - a_{16}) \cdot (x - a_{17}) \cdot (x - a_{18})}{100} \\
I_5(x) &= \frac{300 \cdot x - 1.5 \cdot (x - a_{19}) \cdot (x - a_{20}) \cdot (x - a_{21}) \cdot (x - a_{22})}{100} \qquad (S4) \\
I_6(x) &= \frac{270 - (x - a_{23}) \cdot (x - a_{24}) \cdot (x - a_{25}) \cdot (x - a_{26}) \cdot (x - a_{27})}{100} \\
I_7(x) &= \frac{4600 + (x - a_{28}) \cdot (x - a_{29}) \cdot (x - a_{30}) \cdot (x - a_{31}) \cdot (x - a_{32})}{100} \\
I_8(x) &= \frac{100 + (x - a_{33}) \cdot (x - a_{34}) \cdot (x - a_{35}) \cdot (x - a_{36})}{100} \\
I_9(x) &= \frac{3200 - (x - a_{37}) \cdot (x - a_{38}) \cdot (x - a_{39}) \cdot (x - a_{40})}{100} \\
I_{10}(x) &= \frac{3000 - 160 \cdot x + (x - a_{41}) \cdot (x - a_{42}) \cdot (x - a_{43}) \cdot (x - a_{44})}{100}
\end{aligned}$$

Subsequently, we randomly selected $m$ (from 2 to 7) out of the ten genes as the gene expression trajectories of the regulators ($x_j$ in the following formula) and simulated the steady-state gene expression of its target gene ($y$) by the following model with randomly chosen parameters:

$$\log_2(y) = \sum_j \log_2\left(\lambda_j + (1-\lambda_j)\frac{1}{1+\left(\frac{x_j}{R_j}\right)^{n_j}}\right) + C \quad (S5),$$

where $\log_2(\lambda_j)$ randomly picked uniformly from either (-3,-1) for inhibition or (1,3) for activation, $R_j$ is randomly selected from $(0.1 \cdot \max(x_j), 0.9 \cdot \max(x_j))$, $n_j$ from (2.5, 4.5) and C from (-2,2).

For the benchmarking test, all ten time trajectories and the simulated time trajectory of the target gene $y$ were provided as the input, where the ground truth is the membership of the true regulator genes. Additionally, from the false regulators, we randomly selected $m_d$ genes (ranging from 1 to 9 – $m$) as the decoy regulators in the test. Thus, each testing case has $m$ true regulators, $m_d$ decoys and one target gene. For each gene, we evenly selected 201 time points along the time trajectory and used the corresponding gene expression data as the input data for the tests. In the current benchmark test, we chose $9 \geq m+m_d \geq 4$, and, for each combination of $(m, m_d)$, we conducted a random sampling of the true regulators and decoys for three times. In total, we generated 81 different test cases for benchmarking.

To test the performance of an algorithm in identifying the true regulators under varying levels of noise, we added Gaussian noise of different intensity to the gene expression of the 201 time points from the original gene expression trajectories of the regulators and targets by

$$G'(t) = G(t) + \gamma \cdot N(0,1) \cdot G(t) \quad (S6),$$

where $G(t)$ is the deterministic gene expression at time t, $G'(t)$ is the noisy gene expression at time $t$, $\gamma$ is a scaling factor that adjusts the intensity of the noise. And $N(0,1)$ represents the Gaussian noise with a mean of 0 and a variance of 1. For each noise level, we tested all the 81 combinations of true regulators and decoys. The synthetic benchmark data set is available in our GitHub repository (see **Data Availability**)

**Performance Assessment for benchmarking using the synthetic data**

For benchmarking GRN inference using the synthetic data, we evaluated their performance against ground truth by calculating area under the receiver operating characteristic curve (AUROC) and area under the precision-recall curve (AUPRC). For methods capable of predicting the interactions with signs, we also assessed the AUROC and AUPRC for activating and inhibiting interactions separately.

We obtained the ranked interactions as predicted from each method. Each interaction was weighted using $\log(A - \pi_e + 2)$ in the process of calculating precision and recall, where $A$ is the number of interactions and $\pi_e$ is the rank of the interaction $e$. It is worth noting that NetDes provides a ranking for different combinations of regulators based on reducibility index $I$, instead of a ranking for individual regulators. Thus, in this benchmark test, to compare with other methods, we instead first identified the best predicted combinations for each possible number of regulators and then ranked regulators based on how frequently they showed up in these combinations. Although this evaluation approach did not fully capture NetDes's performance, NetDes still outperformed other methods in predicting regulators in the benchmark test incorporating gene expression noise and decoy genes.

##### Performance Assessment for benchmarking using the BEELINE data

We benchmarked NetDes on the human embryonic stem cell (hESC) and hepatocyte (hHep) scRNA-seq datasets from the BEELINE collection [1]. For each dataset cells were ordered by the pseudotime provided by BEELINE, and genes were ranked according to the BEELINE preprocessing. We defined the initial GRN by the union of non-specific and cell-type-specific ChIP-seq networks, and defined the ground truth as the cell-type-specific ChIP-seq network. Performance for each method was quantified using the area under the precision-recall curve (AUPRC) computed separately for each target gene among the top 1000 HVGs and then averaged across all target genes. We also reported the expected AUPRC of a random predictor (baseline), computed as the average (across target genes) fraction of true regulators (derived from cell-type-specific ChIP-seq) among all candidate regulators (derived from both non-specific and cell-type-specific ChIP-seq), which was 0.63 for hESC and 0.59 for hHep.

##### Performance Assessment for iPSC-to-DE GRN inference using the literature data

NetDes optimizes a GRN model starting from an initial network composed of interactions derived from TF-target databases, including TRRUST (based on text-mining) and Rcistarget (based on TF binding motif data). As a result, the interactions considered by NetDes already have some supporting evidence. However, these interactions may not be direct or context-specific regulation. To further assess the performance of GRN reconstruction on the iPSC-to-DE dataset, we performed an extensive literature survey to identify TF-target relationships within the initial GRN that are supported by published evidence. Details of these literature-supported interactions are provided in **SI Table1**, and they were regarded as the ground truth in this benchmarking.

For each GRN inference method, we constructed core TF networks, each containing a different number of regulatory edges, and only included interactions present in the initial GRN for benchmarking. The only exception for this filtering step was CellOracle, whose candidate interactions are determined by its prebuilt base GRN rather than by the initial GRN. We considered GRNs with 31 to 45 edges (after filtered based on the initial GRN) to ensure that the inferred GRNs remained connected while being substantially smaller than the initial network (which contains 59 edges). For the benchmarking, we evaluated the AUPRC, accuracy, and $F_{0.1}$ scores according to the literature-based evidence. We used $F_{0.1}$ which weights precision ten times more heavily than recall. Omiiting a literature interaction may well be correct, since a minimal, context-specific GRN is expected to drop some interactions which are inactive in state transition, whereas assigning a sign that contradicts the literature is almost certainly an error. The AUPRC values were calculated over GRNs with 31–45 edges, whereas $F_{0.1}$ and accuracy correspond to the highest values achieved among the reconstructed GRNs. In this analysis, a true positive is an interaction both identified in the GRN and supported by the literature with consistent interaction sign, a false positive is an interaction agreed upon by both but with inconsistent signs, and a false negative is an interaction verified in the literature but missed by the GRN.

#### Mapping GRN gene expression states to single cell RNA-seq data

We performed additional model simulations of the inferred GRNs and checked how consistent the GRN gene expression profiles are with the gene expression states from the single cell RNA-seq data. To do this, we selected six time points evenly along the smoothed time trajectories of the GRN TFs as the reference states. Choosing six time points allows for a more comprehensive representation of the dynamic changes in gene expression over time, capturing key cell state transitions and ensuring a robust comparison between the model simulations and the observed gene expression states from the scRNA-seq data. The mapping procedure consists of the following steps.

First, we applied, a mathematical modeling method, Random Circuit Perturbation (RACIPE)[2] , on each inferred GRN, to simulate single cell gene expression profiles. Here, according to the topology of the GRN, RACIPE generates an ensemble of ODE models with randomly selected kinetic parameters and obtained a stable steady state for each model via ODE simulations. Ensemble-based mathematical modeling would allow to capture cell-to-cell variability in a single-cell population and the effects of GRN regulation. For each inferred GRN, RACIPE was applied to generate 10,000 gene expression profiles of the GRN genes.

Second, simulated gene expression data were log-normalized so that each gene's distribution matches that of the smoothed time trajectory. Specifically, the gene expression $u_i$ of gene $i$ in a RACIPE model was log transformed by $log_2(u_i + k_i)$, where $k_i$ is a gene specific constant for the GRN simulation. Afterwards, standardization was applied to the log transformed data. This treatment helps to make RACIPE-simulated data more closely match with the experimental expression data. For each gene $i$, we selected $k_i$ such that $log_2(k_i)$ was evenly sampled between -10 and 5. We then identified the optimal $k_i$ that maximized the overlap between the distribution of the log-normalized RACIPE models and that of log-normalized gene expression from 201 evenly distributed points along the smoothed time trajectory.

Third, to establish the mapping, we calculated the Euclidean distance in high-dimensional gene expression profiles $D_{a,r}$ , which is between a RACIPE model $a$ and one of the six reference states $r$ (after log-normalization). The RACIPE model $a$ was successfully mapped to the reference state $r$, when

$$D_{a,r} \leq D_r \quad \text{(S7)},$$

where $D_r$ is the cutoff distance for the reference state $r$, defined as the bottom $\alpha$% of the Euclidean distances between a random gene expression profile and the expression profile of reference state $r$. The random gene expression profiles were generated from a normal distribution with the same mean and standard deviation of RACIPE simulated gene expression profiles. Here, a different $\alpha$ was chosen for each dataset so that a comparable fraction of RACIPE models was assigned to the reference states, and a larger $\alpha$ was needed for a larger GRN. We chose $\alpha = 5$ for the iPSC-to-DE and EMT networks (with 12 and 8 TFs, respectively); while we chose $\alpha = 10$ for the ERY and DC networks (with 15 and 19 TFs, respectively). If the model $a$ can be mapped to more than one reference states, the model is mapped to the reference state with the minimum $D_{a,r}/D_r$.

We quantified how well RACIPE models are mapped to any of the six reference states by a metric called *weighted mapping percentage*. Here, the mapping of each RACIPE model is weighted by the inverse of its local density of gene expression from all 10,000 RACIPE models. The local density of a model $a$ is defined as

$$\rho_a = 15/\pi {R_a}^{v} \quad \text{(S8)},$$

where $R_a$ is Euclidean distance of the gene expression profiles, considering all data dimension $v$, between the model and the 15th nearest neighbor. In our application, we also calculated the low-dimensional local density, which considered the $R_{a\_low}$ as the distance within the first three principal components (PCs) obtained from PCA, where the dimension $v$ replaced by $v_{low}$ = 3.

Then, the weighted mapping percentage is defined as

$$p_{tot} = \sum_{a} \frac{\sum_{a \epsilon A_r} \frac{1}{\rho_a}}{\sum_{a \epsilon A} \frac{1}{\rho_a}} \quad \text{(S9)},$$

where $A_r$ presents all the RACIPE models mapped to the reference state $r$, and $A$ presents all the RACIPE models. Here, weights $\rho_a$ inversely proportional to model density were used to avoid favoring top-ranked GRNs that achieve high mapping percentages merely by aligning well with only one or two reference states. We incorporated both low-dimensional local density (using the first three principal components) and full-dimensional density when calculating the weighted mapping percentages. It is worth noting that local density computed in the full-dimensional space tends to produce highly skewed distributions. Hence, our analysis is mainly based on low-dimensional mapping results.

To evaluate a GRN's overall performance in capturing various states along a continuous state transition, we defined *the entropy of the mapped state distribution* as

$$H = -\sum_{i=1}^{n_r} p(r_i)\log_2 \mathrm{p}(r_i) \quad \text{(S10)},$$

where $p(r_i)$ represents the percentage of simulated RACIPE models that mapped to a reference state $r_i$, and $n_r$ is the number of the reference states. Higher entropy indicates that a GRN captures states more evenly, whereas lower entropy suggests the GRN is biased toward specific states.

We also used another metric to quantify how gene expression states allowed by a model deviate from the reference states. For those RACIPE models that are not mapped to any reference state, we evaluated how each model deviates from the closest reference state in gene expression. Thus, the *model deviation* was defined as the average normalized distance

$$d_u = \frac{1}{N_u}\sum_{b=1}^{N_u} \min\left(\frac{D_{b,r}}{D_r}\right) \quad (S11),$$

where the summation is over any unmapped RACIPE models $b$, $D_{b,r}$ is the Euclidean distance in high-dimensional gene expression profiles between unmapped RACIPE model $b$ to a reference state $r$. $N_u$ is the total number of unassigned RACIPE models. Similarly, we also computed the average normalized distance $d_\theta$ for random profiles, where the RACIPE simulated gene expression profiles are replaced by the random gene expression profiles previously generated from normal distributions. Finally, we defined the model deviation metric as $\omega = \frac{d_u - 1}{d_\theta - 1}$. A high model deviation metric typically suggests the existence of unmapped gene expression cluster(s) from GRN simulations, in which case the GRN is incapable of fully recapitulating observed gene expression states.

#### Implementation of other GRN reconstruction methods

We benchmarked NetDes against several existing GRN reconstruction method, applying each to the processed expression data of the same core TFs used by NetDes. On the synthetic datasets, we compared NetDes with GENIE3[3], ppcor[4] and SCODE[5]. On the human embryonic stem cell (hESC) and hepatocyte (hHep) scRNA-seq data from BEELINE[1], we additionally included the deep-learning-based methods, DeepSEM[6] and DeepRIG[7]. On the four experimental cell-state-transition datasets (iPSC-to-DE, EMT, ERY and DC), we compared NetDes with GENIE3, ppcor, SCODE, SINCERITIES[8] and CellOracle[9]; for each dataset, the edges inferred by each method were restricted to the same core TFs as NetDes before downstream simulation and evaluation. The implementation of these methods is outlined as follows.

The ppcor[4] method calculates partial and semi-partial correlation coefficients between regulators and target genes. In our testing, we used semi-partial correlation for obtaining the interactions with sign. For synthetic data, we opted for the Kendall method to compute the correlation matrix, instead of using the default Pearson correlation coefficients due to errors associated with zero eigenvalues. For scRNA-seq data, we applied ppcor to the processed data, and GRN interactions were ranked based on the absolute value of Pearson (or Kendall) correlation coefficients.

GENIE3[3] employs a tree-based model to predict the interactions between genes. We utilized the default setting of GENIE3 for its application to both the synthetic data and processed scRNA-seq data. To specify the activation and inhibition nature of each regulator-target regulation, we assigned the sign of Pearson correlation coefficients to the interaction. Additionally, the regulatory edges were ranked by the interaction weights inferred by GENIE3.

SCODE[5] utilizes a linear ODE method to fit the GRN dynamics. We used default setting and running 100 iterations of optimization for both the synthetic data and processed scRNA-seq data. Regulatory edges were ranked by the absolute value of correlation coefficient defined by SCODE.

CellOracle[9] involves learning from chromatin accessibility data and gene expression data. We applied CellOracle to the raw scRNA-seq data of the iPSC-to-DE, EMT, ERY and DC datasets using its own prebuilt GRN, derived from motif scanning with gimmemotifs (hg19 for the human iPSC-to-DE, EMT and DC datasets; mm10 for the mouse ERY datasets). For every dataset we report two cluster settings: all cells in a single cluster, giving one fit per edge as for the other methods; and one network per cell state (EMT, treatment time points; ERY, annotated cell types; DC, sampling points; iPSC-to-DE, day labels), keeping each edge's highest score across clusters. The inferred TF–TF edges were assigned activation/inhibition by the sign of the regression coefficient and ranked by the absolute value of the coefficient.

SINCERITIES[8] analyzes gene expression changes over time with the Kolmogorov-Smirnov statistic. In the implementation, we chose the ridge regression from four different regularization regression strategies to find the gene interactions with sign using the processed scRNA-seq data. The regulatory edges were ranked based on the absolute values of their weights, as determined by SINCERITIES, which reflects both the strength and direction of interactions. This ranking was used to construct GRNs of different sizes.

DeepSEM[6] employs a deep structural-equation model, implemented as a β-variational autoencoder, to infer directed regulatory interactions between genes. We applied DeepSEM in an unsupervised manner, using as input the expression of the genes in the initial GRN. We ran the non-cell-type-specific GRN inference task in BEELINE with the default setting for 90 epochs as a single run, and the regulatory edges were ranked by the absolute values of the weights inferred by DeepSEM.

DeepRIG[7] utilizes a graph autoencoder to infer regulatory interactions from a correlation-based co-expression network derived from gene expression. In contrast to the other methods, DeepRIG requires a network of known interactions for training; In BEELINE benchmarking, we provided the union of cell-type-specific and non-specific ChIP-seq network as the training labels and applied cross-validation using the default settings. The regulatory edges were ranked based on the interaction probabilities predicted by DeepRIG.

#### Supplementary Table

##### Supplementary Table 1. Literature evidence for TF-target interactions from the initial GRN.

For every regulatory interaction from the initial GRN, the column named "Interaction type" indicates whether the transcriptional regulation is activating or inhibitory (blank if unknown). The column named "Reference" describes literature evidence and their references. An "NA" in this column represents that no supporting publication was identified.

| Source | Target | Interaction type | Reference |
|---|---|---|---|
| E2F4 | TWIST2 | | NA |
| FOS | TWIST2 | | NA |
| LEF1 | TWIST2 | | NA |
| MYCN | TWIST2 | | NA |
| RARA | TWIST2 | | NA |
| GATA3 | TWIST2 | | NA |
| LEF1 | HES1 | | NA |
| LEF1 | RARA | | NA |
| LEF1 | ZEB1 | Activation | (indirect)The EMT induced by LEF-1 is associated with an increase in the mRNA levels of ZEB1[10]. (https://doi.org/10.1016/j.bbrc.2013.11.031) Zeb1 is indirect recruited to regulatory regions by Lef1 results in gene activation[11] . (https://doi.org/10.15252/embj.201797115) |
| LEF1 | GATA3 | Inhibition | LEF-1 suppresses GATA-3 DNA-binding activity and Th2 cytokine production[12]. (https://doi.org/10.1111/j.1365-2567.2008.02854.x) |
| FOS | RARA | | NA |
| E2F4 | GATA3 | | NA |
| E2F4 | HES1 | | NA |
| E2F4 | LEF1 | | NA |
| E2F4 | RARA | | NA |
| E2F4 | ZEB1 | | NA |
| SOX2 | GATA3 | | NA |
| SOX2 | HES1 | | NA |
| SOX2 | LEF1 | Inhibition | Sox2 acts as a repressor to directly modulate Wnt-responsive transcription of the Lef-1 gene promoter[13]. (https://doi.org/10.1152/ajplung.00157.2013) |
| SOX2 | ZEB1 | Inhibition | (indirect) SOX2 and OCT4 transcriptionally regulate miR200, which in turn represses ZEB genes[14]. (https://doi.org/10.1073/pnas.1212769110) |
| SOX2 | RARA | Inhibition | (indirect) SOX2 knockdown in HF2303 GBM Cells increases RARA expression by 2.2 fold change[15]. (https://doi.org/10.1016/j.neo.2014.03.006) |
| ZEB1 | RARA | Activation | (indirect) Knockdown of ZEB1 suppresses the RARA-mediated EMT phenotype[16]. (https://doi:10.1016/j.molonc.2014.09.005) |
| POU5F1 | LEF1 | Activation | Oct4 induces EMT through LEF1/β-catenin dependent WNT signaling pathway in hepatocellular carcinoma[17]. (https://doi.org/10.3892/ol.2017.5788) |
| POU5F1 | RARA | | NA |
| POU5F1 | GATA3 | | NA |
| POU5F1 | ZEB1 | Inhibition | (indirect) SOX2 and OCT4 transcriptionally regulate miR200, which in turn represses ZEB genes[14]. (https://doi.org/10.1073/pnas.1212769110) |
| ZEB1 | LEF1 | | NA |

| Source | Target | Interaction type | Reference |
| --- | --- | --- | --- |
| GATA3 | RARA | Activation | (indirect) RARα can be recruited to GATA binding sites by protein interactions[18]. (https://doi.org/10.1128/MCB.24.15.6824-6836.2004) |
| FOS | ZEB1 | | NA |
| GATA3 | HES1 | Activation | Chip-seq shows that GATA3 binds to the promoter of HES1[19]. (https://doi.org/10.1016/j.stem.2020.03.005); HES1 is upregulated following GATA3 overexpression[20] (https://doi.org/10.1038/ncomms11171) |
| FOS | GATA3 | | NA |
| GATA3 | LEF1 | | NA |
| POU5F1 | HES1 | | NA |
| FOS | LEF1 | | NA |
| KLF8 | RARA | | NA |
| RARA | GATA3 | Activation | RARA can be recruited to GATA3 binding sites to influence GATA3 activity at its target genes[18]. (https://doi.org/10.1128/MCB.24.15.6824-6836.2004) |
| MYCN | RARA | Inhibition | MYCN inhibits normal RA-mediated neuronal differentiation[12] (https://doi.org/10.1186/s13073-017-0407-3) |
| MYCN | ZEB1 | | NA |
| MYCN | HES1 | Activation | MYCN binds to the HES1 promoter and exhibits transcriptional activity[21]. (PMID: 31598396) |
| KLF8 | GATA3 | | NA |
| KLF8 | HES1 | | NA |
| KLF8 | LEF1 | | NA |
| POU5F1 | MYCN | Activation | MYCN and its cis-antisense gene, NCYM, form a positive feedback loop with OCT4[22]. (https://doi.org/10.1111/cas.12677) |
| POU5F1 | SOX2 | Activation | Oct4 and Sox2 bind directly to the composite sox-oct elements in both Pou5f1 and Sox2 in living mouse and human ESCs[23]. (https://doi.org/10.1128/MCB.25.14.6031-6046.2005) |
| ZEB1 | KLF8 | | NA |
| GATA3 | MYCN | | NA |
| GATA3 | SOX2 | Inhibition | (indirect) MTA3 is recruited by GATA3 to repress SOX2OT transcription and then SOX2[24]. (https://doi.org/10.1016/j.isci.2019.11.009) |
| GATA3 | KLF8 | | NA |
| KLF8 | MYCN | | NA |
| MYCN | KLF8 | Activation | MYCN up-regulates PTK2; KLF8 is positively regulated by PTK2[25]. (https://doi.org/10.3390/molecules28031141) |
| E2F4 | FOS | | NA |
| GATA3 | FOS | Activation | GATA3 binds to the promoter region of the FOS gene and activates FOS transcription[26]. (https://doi.org/10.1038/s41419-023-05888-9) |
| GATA3 | E2F4 | | NA |
| RARA | FOS | Inhibition | RARA can directly interact with and inhibit AP-1 transcriptional activity, which includes FOS as a key component[27]. (https://doi.org/10.1210/mend.13.2.0237) |
| MYCN | FOS | | NA |
| HES1 | FOS | | NA |
| GATA3 | ZEB1 | Inhibition | Wild-type GATA3 transcriptionally suppresses ZEB1[28]. (https://doi.org/10.2147/CMAR.S147973) |
| TWIST2 | FOS | | NA |
| ZEB1 | POU5F1 | | NA |

**Supplementary Table 2. Model parameters of the optimized ODEs for the gene circuit governing ICM-TE transition.**

The rate equations for this model are provided in **Eq. S2** in the Methods section. Each gene $i$ corresponds to Oct4 ($x_1$), Cdx2 ($x_2$), Esrrb ($x_3$) and Fgf ($x_4$). $x(0)$ denotes the initial expression levels of the simulated trajectory (ICM-like state), and x(T) denotes the model-predicted expression levels at the final time point T (TE-like state). All parameters are given in arbitrary units.

| **Model parameters** | $x_1$ | $x_2$ | $x_3$ | $x_4$ |
|---|---|---|---|---|
| $x(0)$, **Initial condition** | 3.775 | 72.674 | 119.907 | 150.0 |
| $x(T)$, $x$ **at the final time point** | 3.006 | 97.001 | 114.88 | 25 |
| $g_i$, **Production rate** | 1.5 | 17.0 | 75.0 | 20.0 |
| $k_i$, **Degradation rate** | 0.1 | 0.3 | 0.2 | 0.1 |
| $\lambda_{1\to i}$, **Maximum Fold-change** | 1.0 | 0.04 | 1.0 | 1.0 |
| $\lambda_{2\to i}$, **Maximum Fold-change** | 0.05 | 1.0 | 0.3 | 1.0 |
| $\lambda_{3\to i}$, **Maximum Fold-change** | 5.0 | 1.0 | 1.0 | 1.0 |
| $\lambda_{4\to i}$, **Maximum Fold-change** | 1.0 | 4.0 | 1.0 | 1.0 |
| $R_{1\to i}$, **Hill threshold** | 1.0 | 4.0 | 1.0 | 1.0 |
| $R_{2\to i}$, **Hill threshold** | 10.0 | 1.0 | 30.0 | 1.0 |
| $R_{3\to i}$, **Hill threshold** | 10.0 | 1.0 | 1.0 | 1.0 |
| $R_{4\to i}$, **Hill threshold** | 1.0 | 120.0 | 1.0 | 1.0 |
| $n_{1\to i}$, **Hill Coefficient** | 1.0 | 4.0 | 1.0 | 1.0 |
| $n_{2\to i}$, **Hill Coefficient** | 4.0 | 1.0 | 4.0 | 1.0 |
| $n_{3\to i}$, **Hill Coefficient** | 4.0 | 1.0 | 1.0 | 1.0 |
| $n_{4\to i}$, **Hill Coefficient** | 1.0 | 2.0 | 1.0 | 1.0 |
| $T$, **Total simulation Time** | 1000 | 1000 | 1000 | 1000 |

### Supplementary Figures

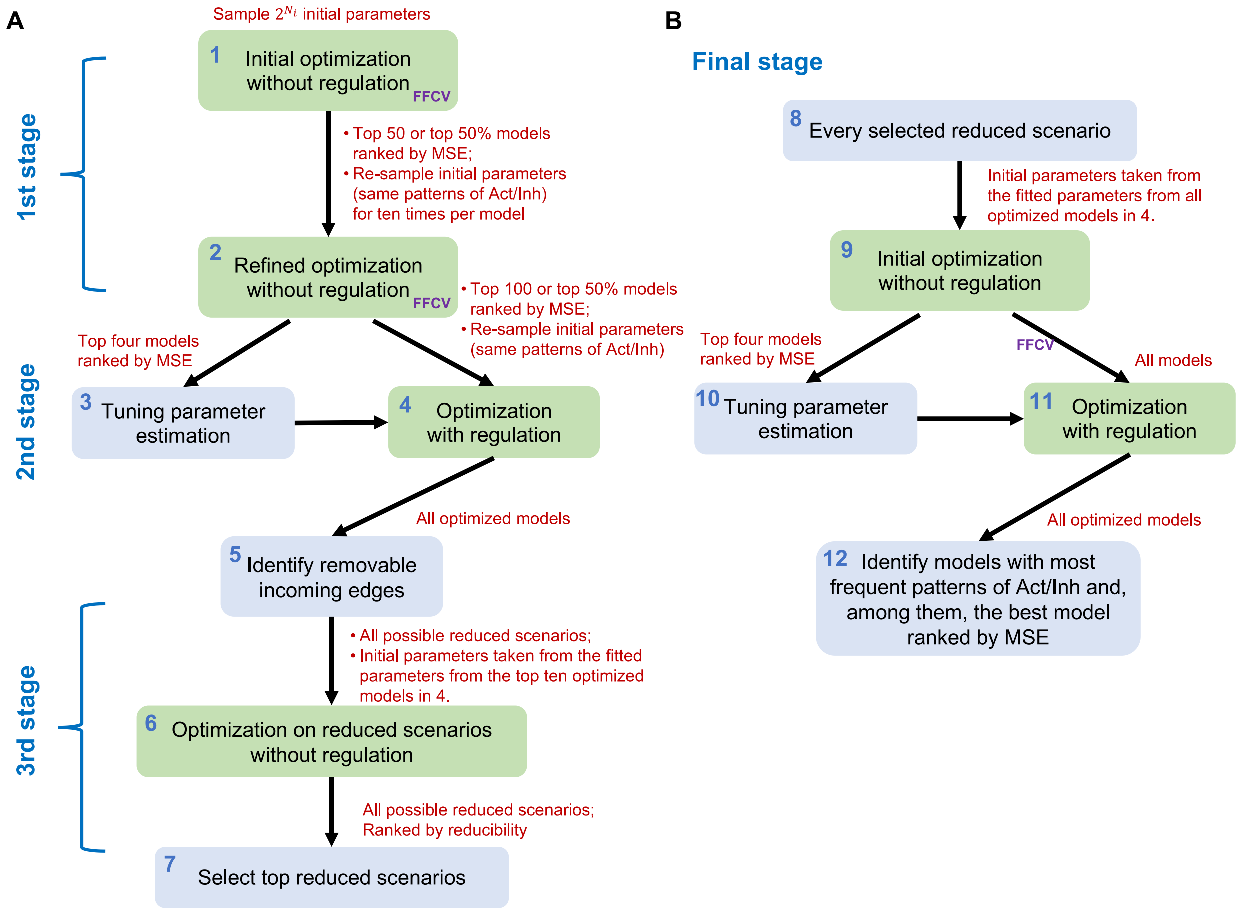

**Fig.S1. Flowchart diagram for the ODE parameter optimization procedure in NetDes.** The whole process consists of four stages of optimization. **(A)** The first, second and third stages of optimization. **(B)** The final stage of optimization.

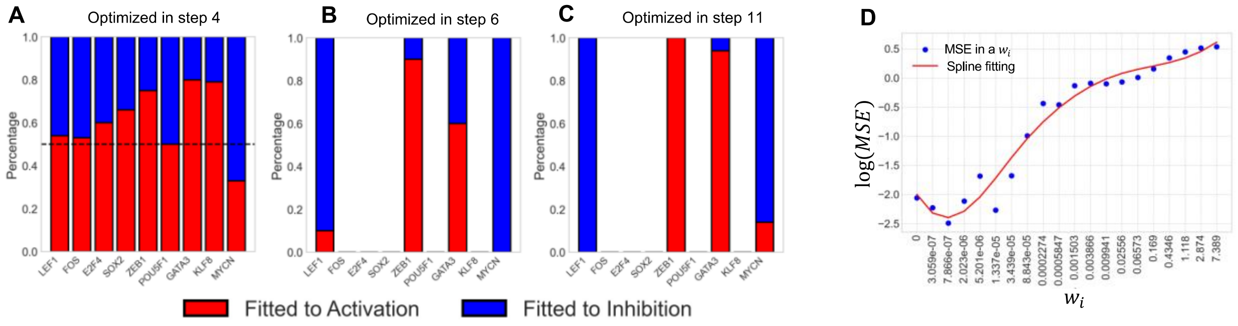

**Fig.S2. Additional model optimization results for the application to time-series scRNA-seq data for iPSC-to-DE differentiation.** Panels **A-C** show, for each putative regulator of RARA, the percentages of optimized models inferred as activating (red) versus inhibiting (blue). These plots illustrate the outcomes at successive steps of the model optimization protocol (see **Fig. S1**): deep sampling during step 4 (A), edge removal in step 6 (B), and incorporate regulation terms in step 11(C). In later steps, the distributions are increasingly dominated by either activating or inhibiting interaction, suggesting more robust and convergent fitting outcomes. **(D)** Example of tuning parameter estimation. The scatter plot shows log(MSE) values across different values of the tuning parameter $w_i$ (blue dots). The red curve indicates a spline fit to the data. The final value of $w_i$ was selected as the one corresponding to the minimum log(MSE) from the curve.

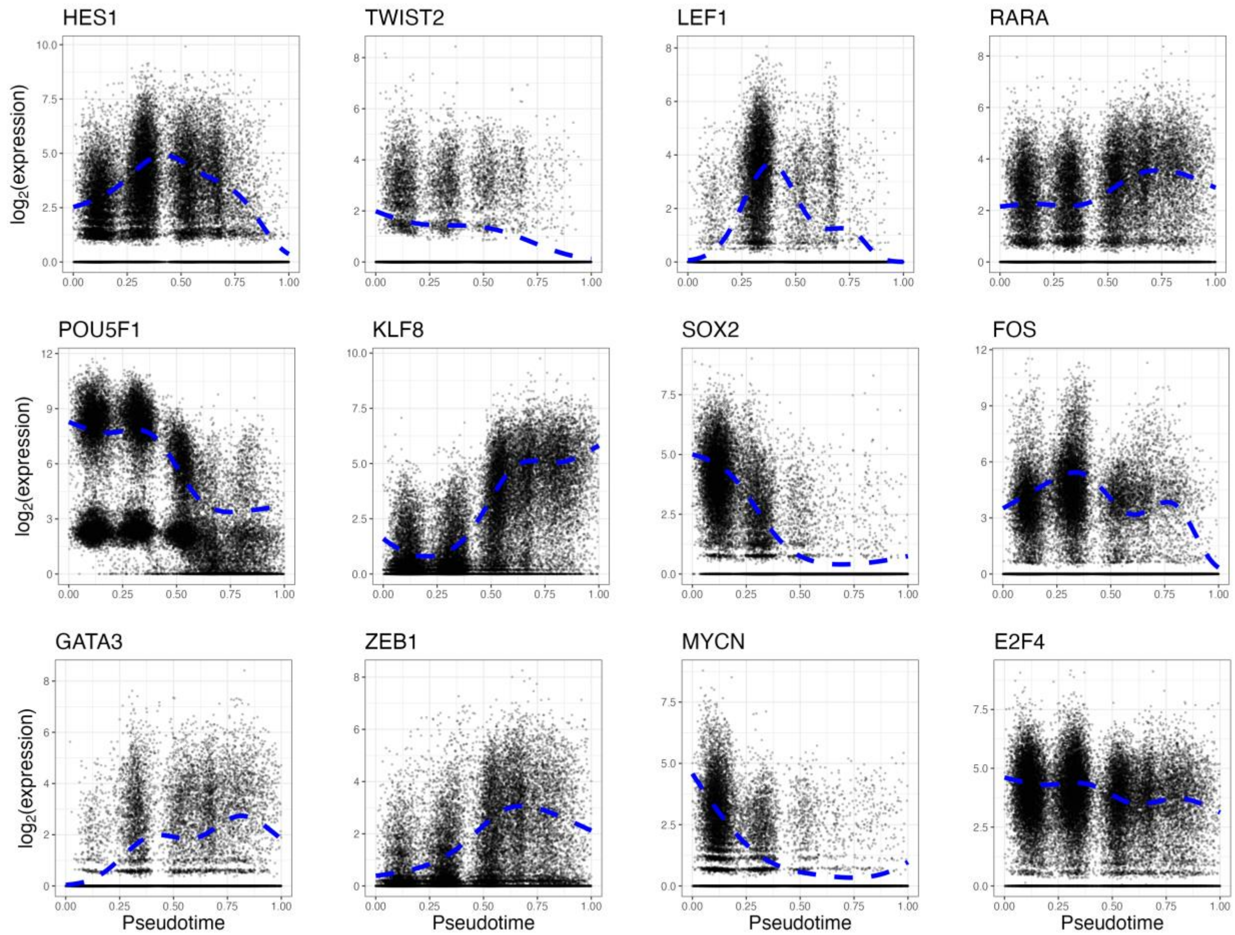

**Fig.S3. Examples of smoothed gene expression trajectories.** Examples of smoothed gene expression trajectories along the pseudotime (in blue dashed lines) for selected transcription factors (TFs), as inferred by PseudotimeDE using single cell expression levels (black dots).

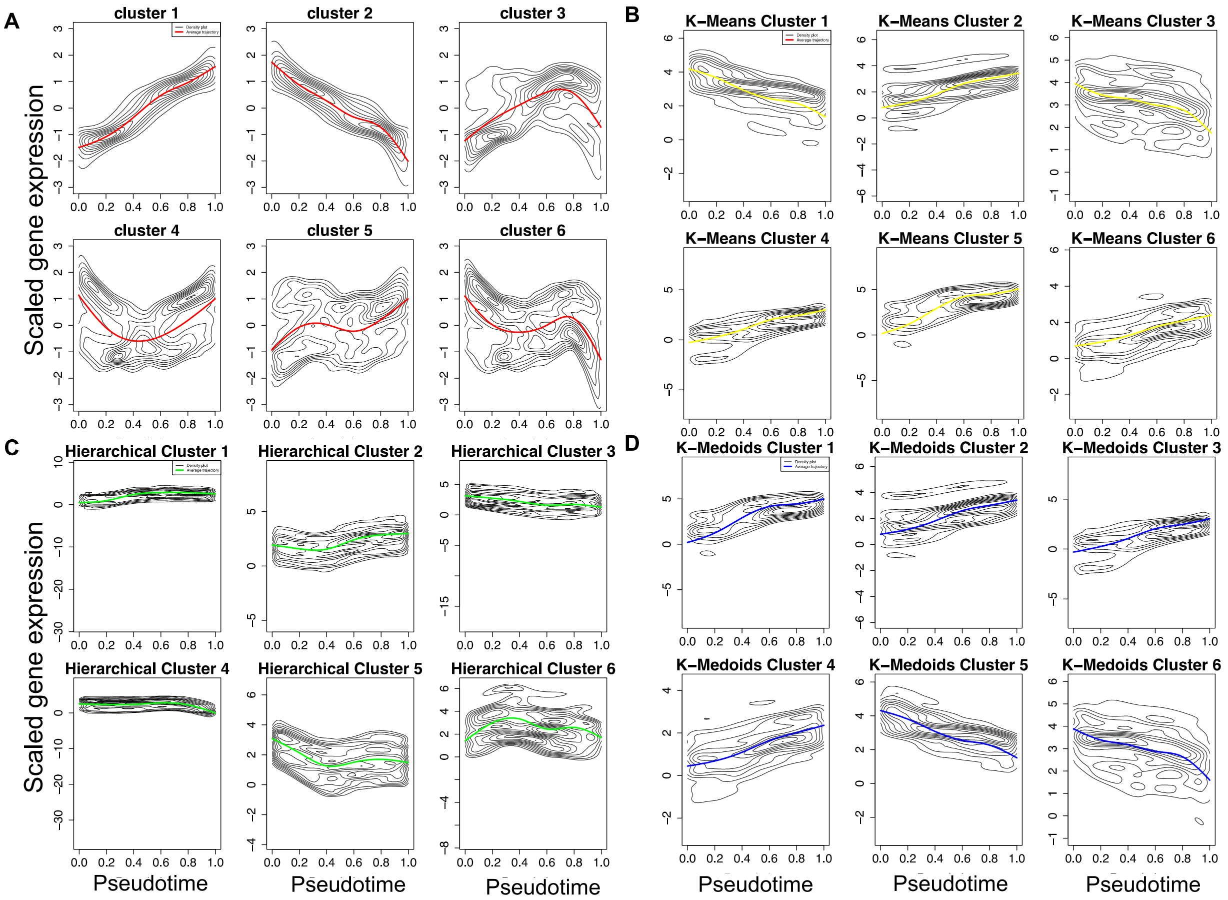

**Fig.S4. Comparison of clustering algorithms on gene expression trajectories.** Smoothed gene expression trajectories were grouped into six clusters by four different algorithms. In all panels, gray contour lines indicate the density of individual trajectories, and the colored curve represents the average trajectory of each cluster. **(A)** NetDes (red), **(B)** K-means (yellow), **(C)** hierarchical clustering (green), and **(D)** K-medoids (blue). Compared to the other methods, NetDes clustering yields more distinct and informative patterns in the time trajectories.

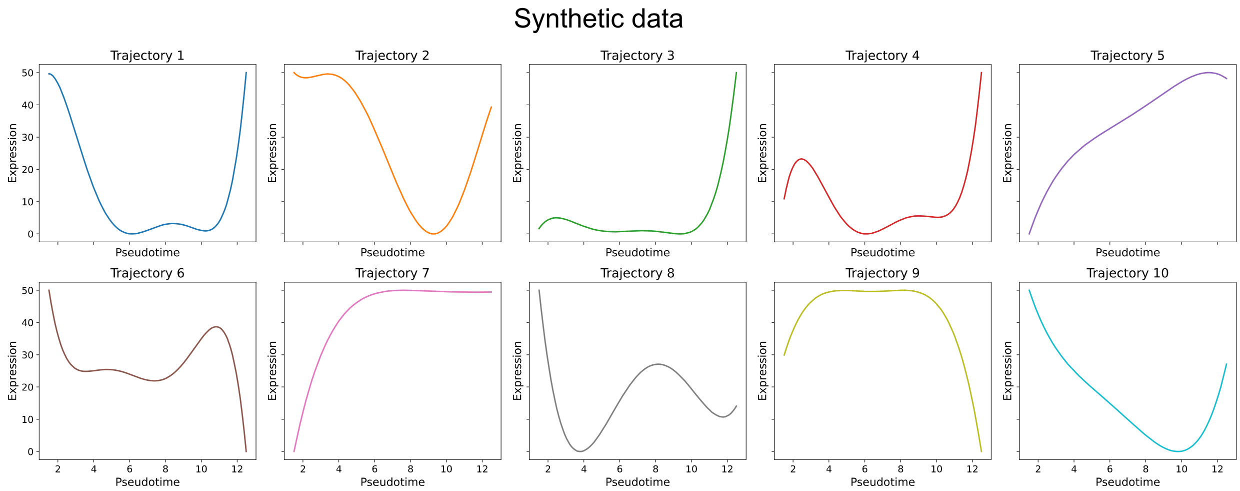

**Fig.S5. Ten time trajectories for synthetic benchmarking.** These trajectories were generated as described in **Eq. S4** and represent a diverse range of dynamic behaviors.

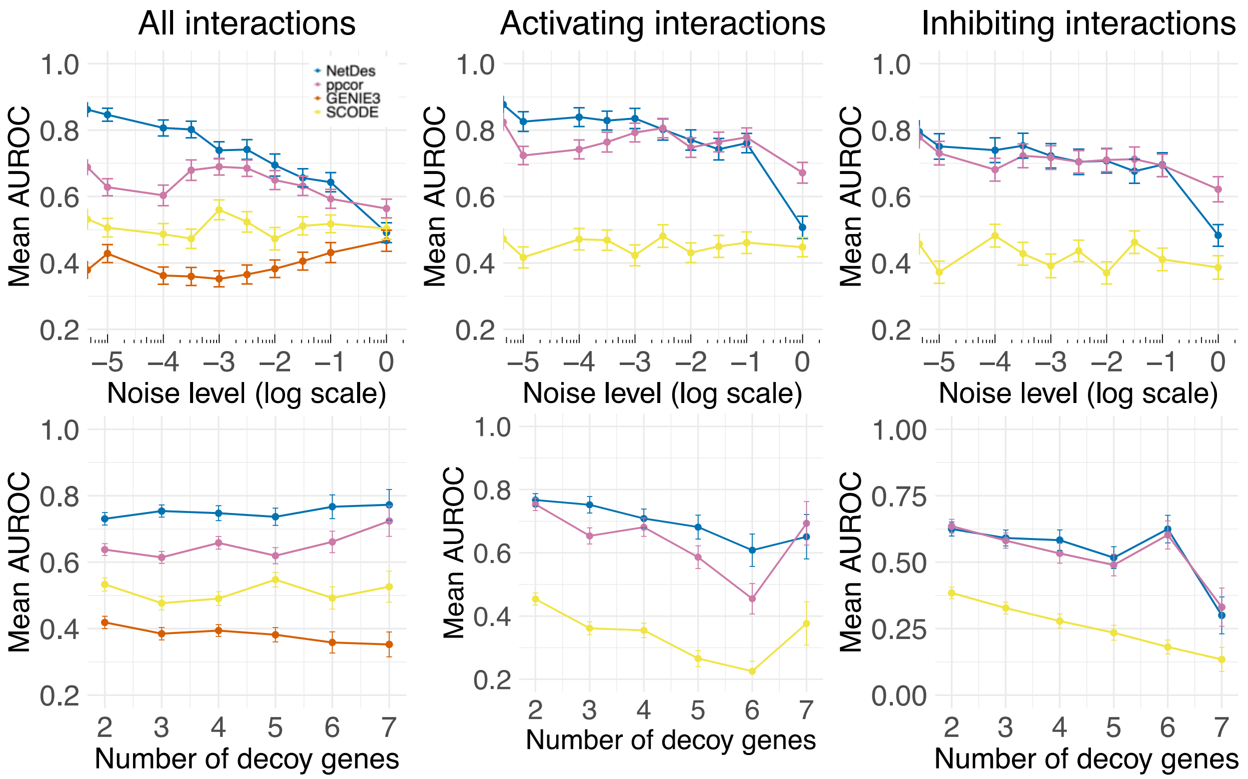

**Fig.S6. AUROC results for benchmarking NetDes and other inference methods on the synthetic dataset.** Top panels show the mean AUROC across different noise levels (in log scale) for all interactions (*i.e.*, both activating and inhibiting), activating interactions, and inhibiting interactions. Bottom panels show the mean AUROC across different numbers of decoy genes.

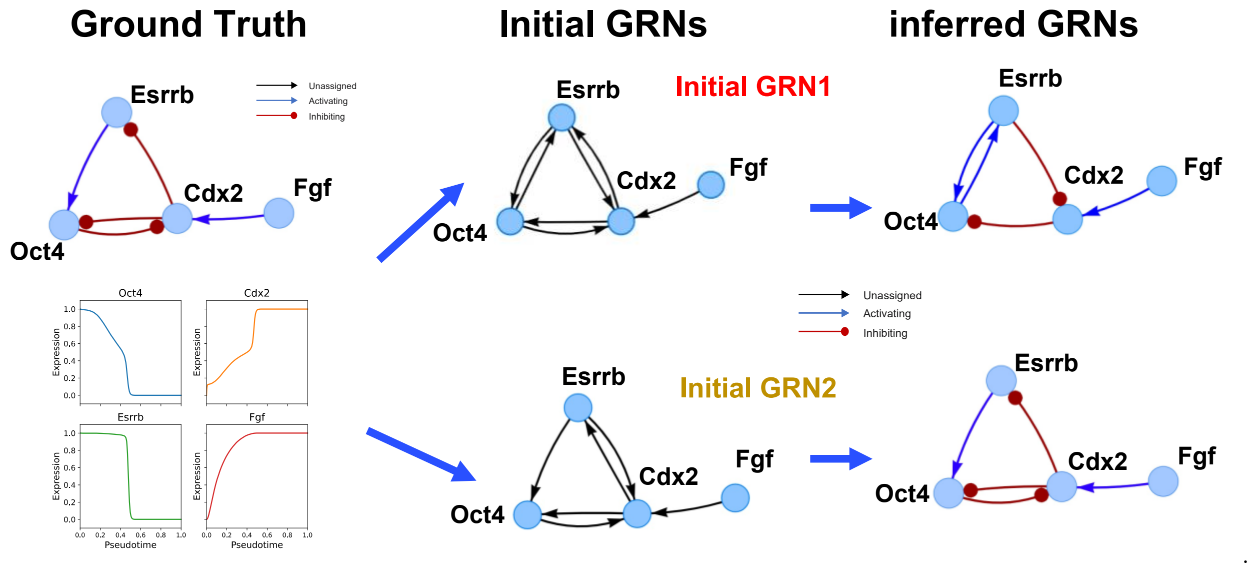

**Fig.S7. NetDes inference of GRN driving cell state transition from inner cell mass (ICM) to trophectoderm (TE).** The top-left panel shows the topology of the ground-truth GRN, where Fgf acts as an input node driving a three-gene circuit consisting of Oct4, Cdx2 and Esrrb. Two distinct initial GRNs (Initial GRN1 and Initial GRN2, middle panels) were provided to NetDes along with synthetic time-series expression data (bottom-left panel). The rightmost panels show the topologies of the GRNs inferred by NetDes.

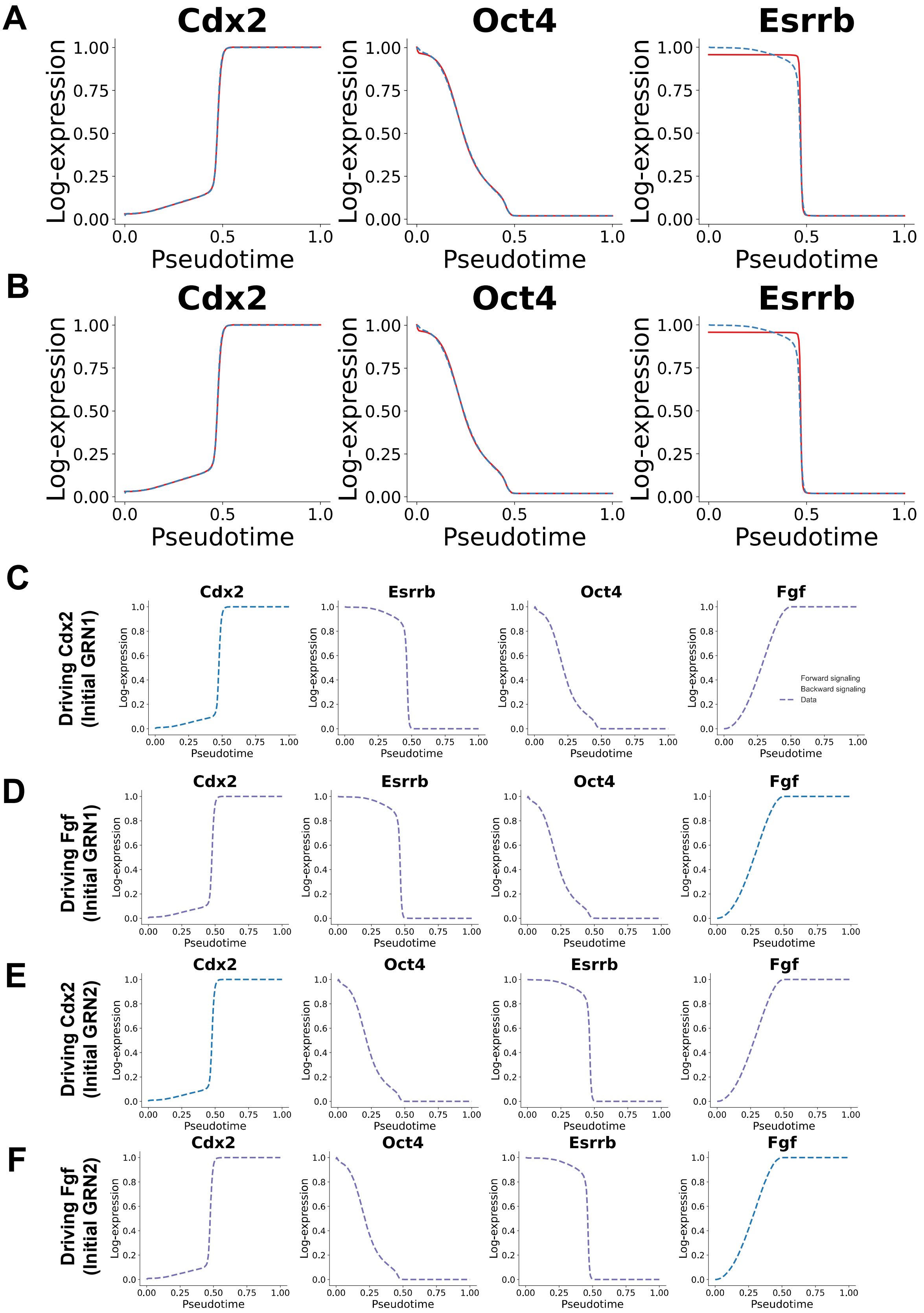

A
Cdx2
Oct4
Esrrb
Log-expression
Pseudotime
B
Cdx2
Oct4
Esrrb
Log-expression
Pseudotime
C
Driving Cdx2 (Initial GRN1)
Cdx2
Esrrb
Oct4
Fgf
Forward signaling
Backward signaling
Data
D
Driving Fgf (Initial GRN1)
Cdx2
Esrrb
Oct4
Fgf
E
Driving Cdx2 (Initial GRN2)
Cdx2
Oct4
Esrrb
Fgf
F
Driving Fgf (Initial GRN2)
Cdx2
Oct4
Esrrb
Fgf

**Fig.S8. GRN simulations for NetDes inferred GRNs for the ICM-to-TE state transition. (A-B)** Comparison of gene expression trajectories for Oct4, Esrrb and Cdx2 between the ground-truth synthetic data (blue dashed lines) and the simulated trajectories from the NetDes model (red solid lines) for the two inferred GRNs. Panel A corresponds to the upper-right GRN in **Fig.S5**, and panel B to the lower-right GRN. **(C-D)** Simulated gene expression trajectories when the first inferred GRNs was driven by Cdx2 (C) and Egf (D). Solid lines represent the simulated trajectories when the GRN was driven by the input signals along the forward (orange) and backward (green) directions. **(E-F)** Corresponding simulation results for the second inferred GRN.

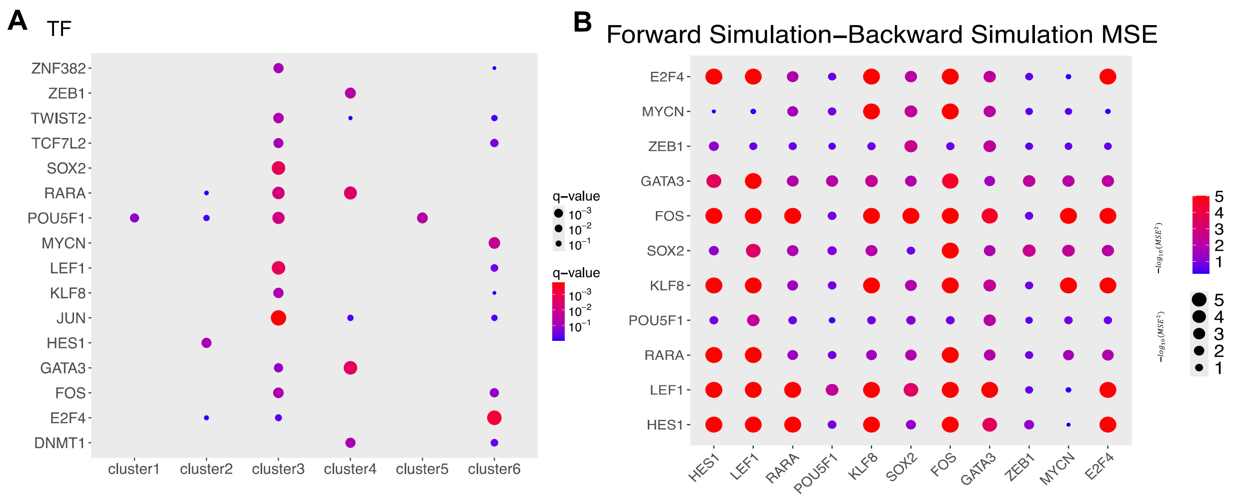

**Fig.S9. Core TF enrichment and TF driving analysis for the inferred iPSC-to-DE GRN. (A)** Gene enrichment analysis using literature-based TF-target gene sets identify significant TFs for each gene cluster (q-value < 0.1). **(B)** The diagram shows mean squared error (MSE) between the forward and backward simulated gene expression trajectories of the optimized iPSC-to-DE GRN, driven by one (diagonal) or two (off-diagonal) TFs; dot size and color represent the MSE.

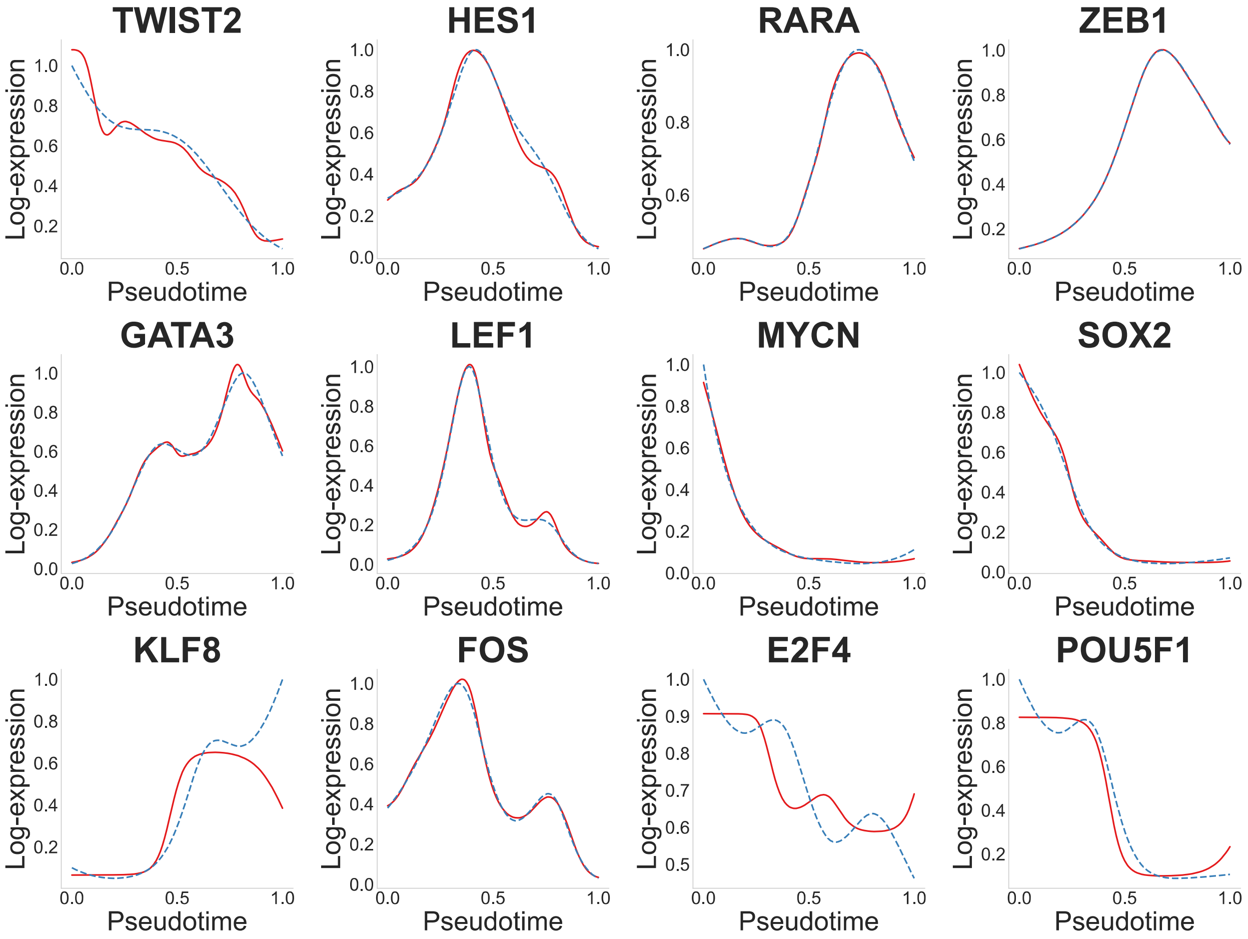

**Fig.S10. Comparison between experimental and fitted gene expression trajectories for each TF.** Dashed blue curves show the log-transformed, smoothed gene expression trajectories from the scRNA-seq data. Solid red curves show the simulated trajectories generated by the optimized model, using regulators' expression trajectories derived from the same scRNA-seq data.

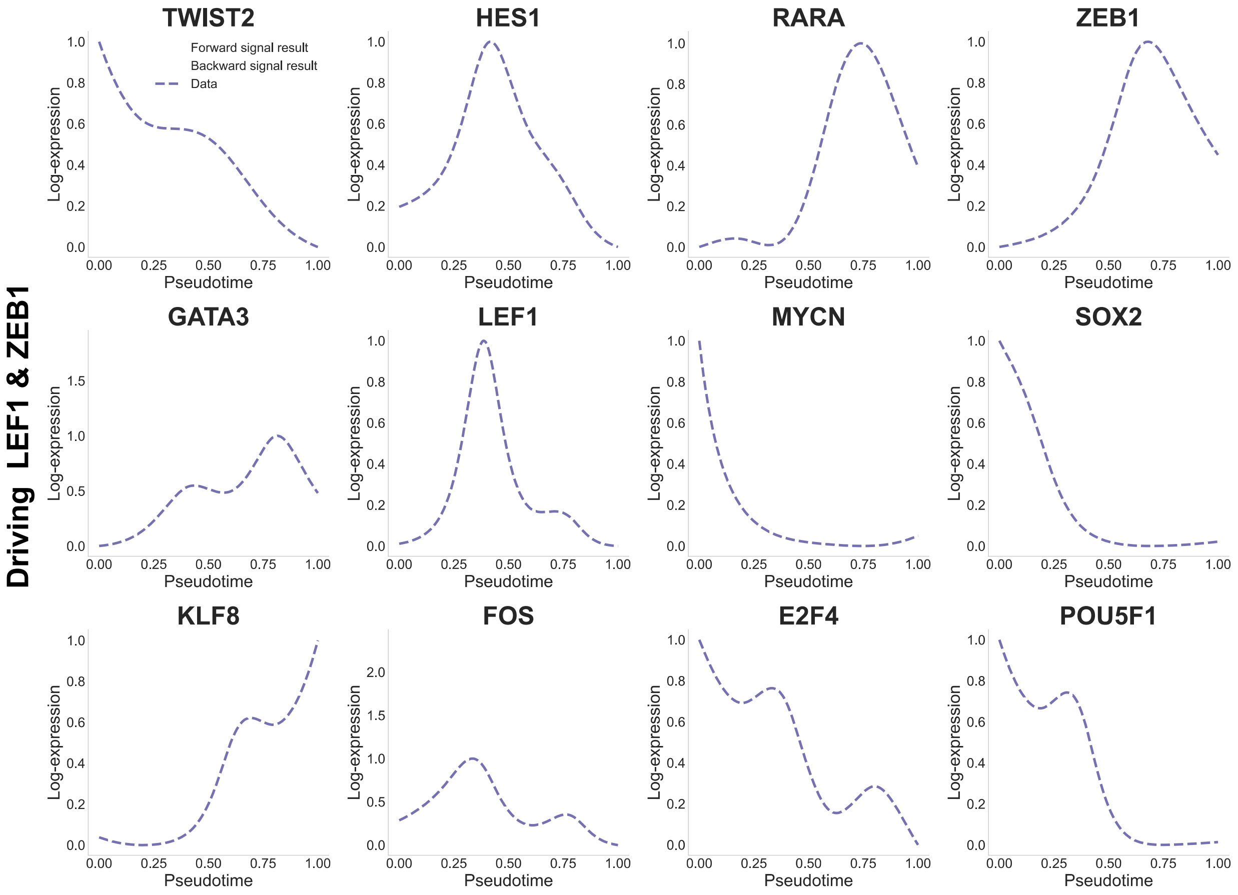

**Fig.S11. Comparison of simulated gene expression trajectories when the optimized GRN was driven by both LEF1 & ZEB1.** The plot is related to **Fig.3E**, with the trajectories for all genes presented here. Each plot shows the smoothed gene expression trajectories along the pseudotime (in blue dashed line), the simulated trajectories for the GRN driven by forward signaling (orange solid line) and backward signaling (green solid line).

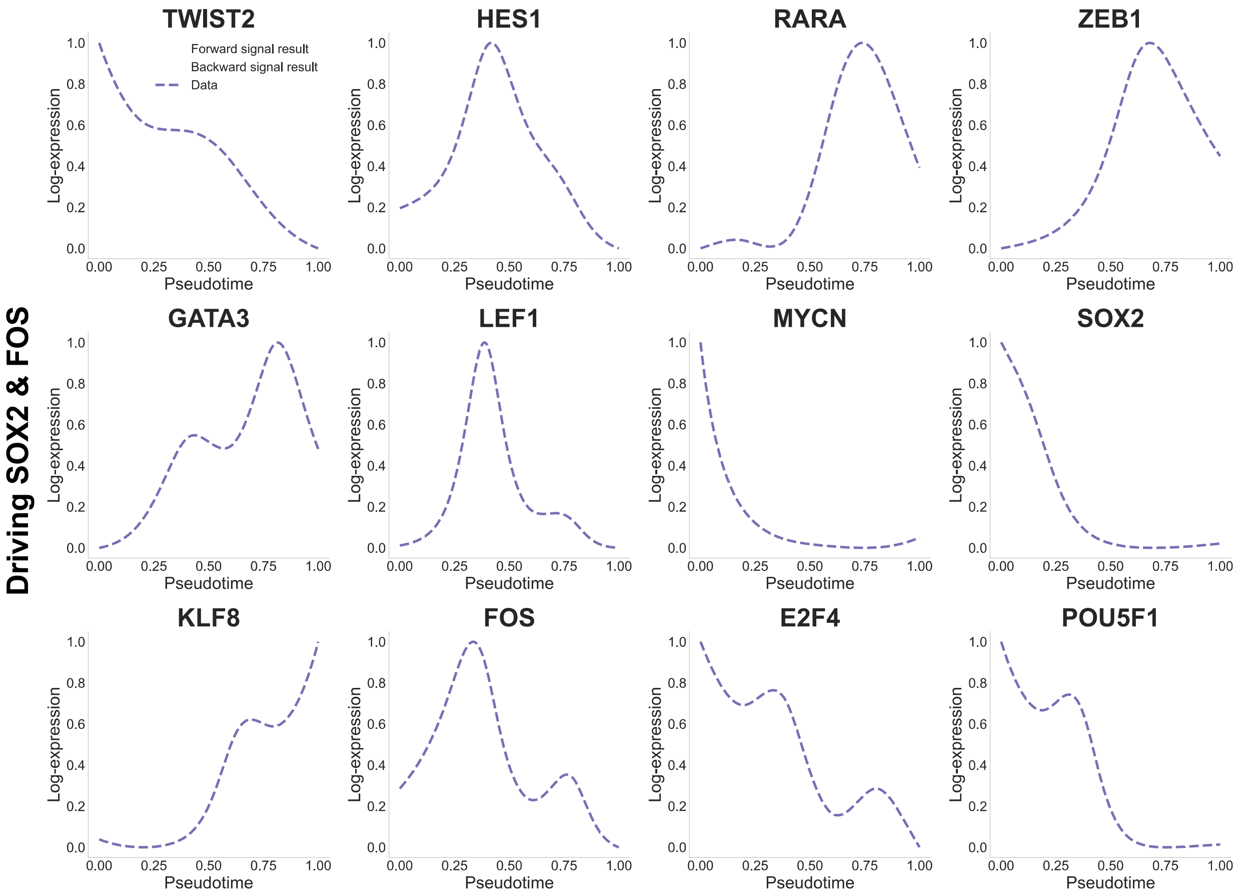

**Fig.S12. Comparison of simulated gene expression trajectories when the optimized GRN was driven by both SOX2 & FOS.** The plot is related to **Fig.3E**, with the trajectories for all genes presented here. Each plot shows the smoothed gene expression trajectories along the pseudotime (in blue dashed line), the simulated trajectories for the GRN driven by forward signaling (orange solid line) and backward signaling (green solid line).

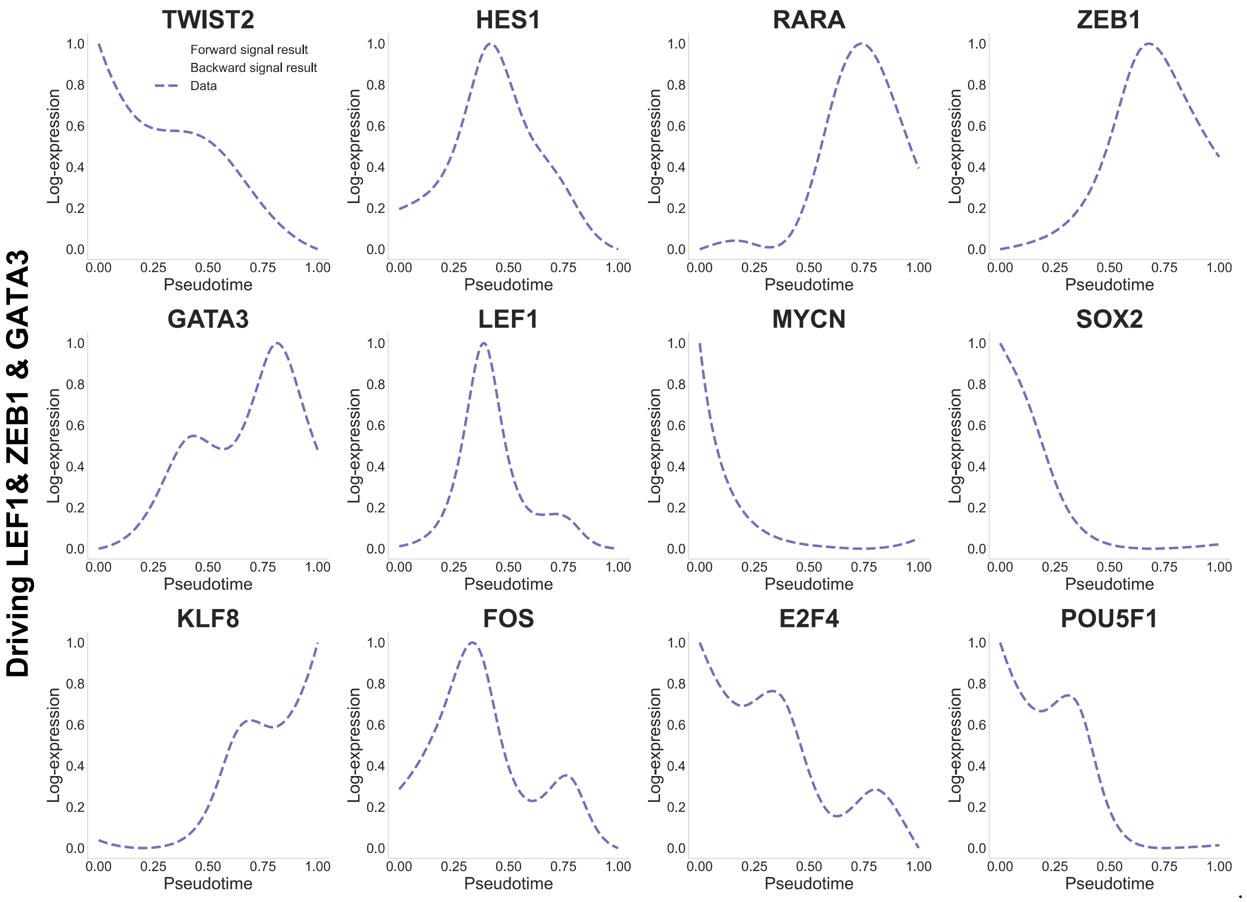

**Fig.S13. Comparison of simulated gene expression trajectories when the optimized GRN was driven by LEF1, ZEB1 and GATA3.** The plot is related to **Fig.3E**, with the trajectories for all genes presented here. Each plot shows the smoothed gene expression trajectories along the pseudotime (in blue dashed line), the simulated trajectories for the GRN driven by forward signaling (orange solid line) and backward signaling (green solid line).

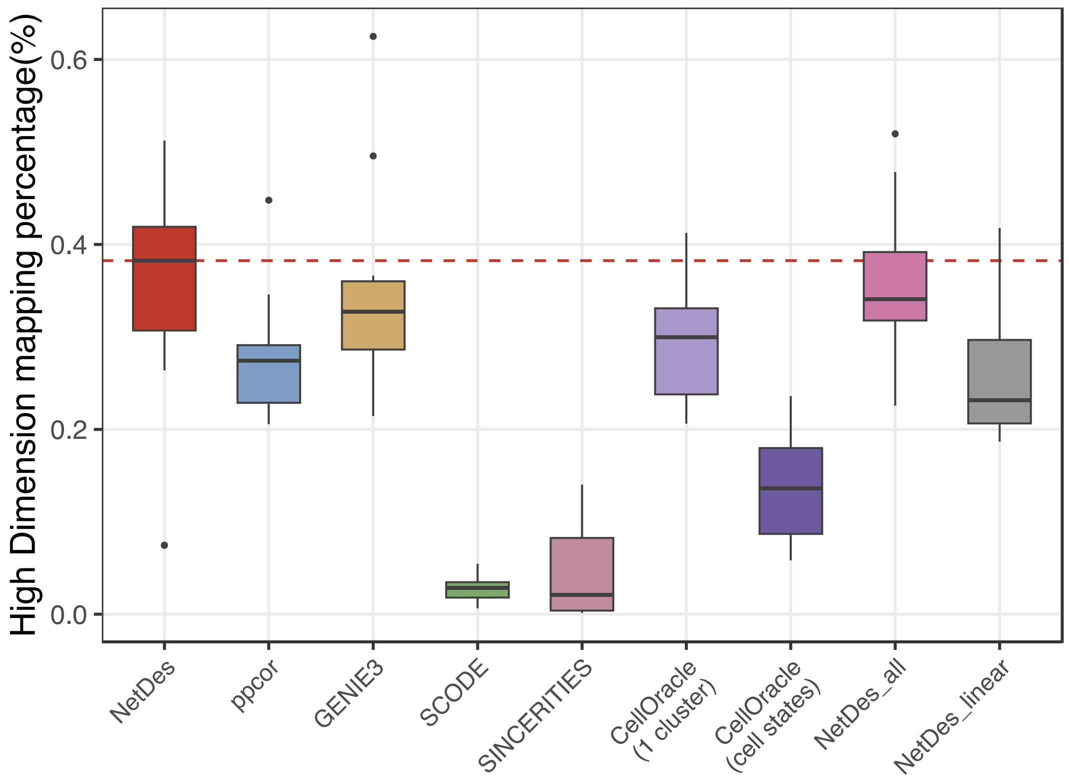

**Fig.S14. Mapping percentages in full-dimensional expression space without providing initial GRN.** (related to **Fig.4**) Box plots showing the full dimension mapping percentages of simulated gene expression profiles to the reference profiles for each inference method.

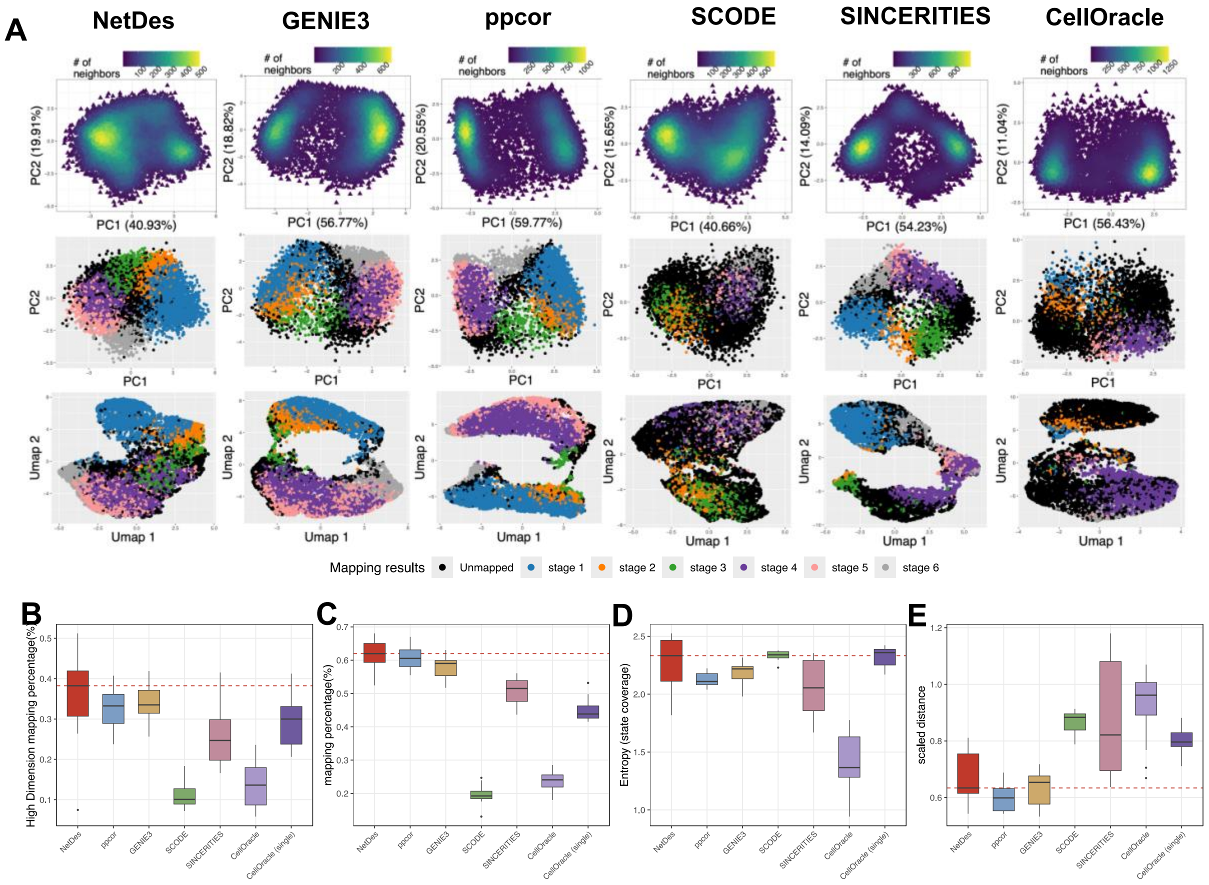

**Fig.S15. Capability of GRN inference in capturing single cell gene expression states during iPSC-to-DE differentiation.** (related to **Fig.5**) In this case, the same initial network as described in **Table.S1** was provided for each GRN inference method. For method other than NetDes, inferred regulatory interactions were only kept if they also present in the initial network. **(A)** Simulated gene expression profiles for the inferred GRNs with the highest mapping percentage for each method. Each set of subplots shows the density plot of simulator gene expression projected onto the first two principal components (top), the scatter plot of gene expression colored by their mapped reference states on the PCA space (middle), and the same scatter plot but projected onto the first two UMAP dimensions (bottom). **(B)** Box plots showing the full dimension mapping percentages of simulated gene expression profiles to the reference profiles for each inference method. **(C)** Box plots showing the low dimension mapping percentages (PC1 – PC3) of simulated gene expression profiles to the reference profiles for each inference method **(D)** Box plots showing the entropy metric that quantifies the disproportion of mapped states. **(E)** Bar plots with scatter points showing the scaled distances in gene expression of unmapped simulated gene expression profiles to the nearest reference profile.

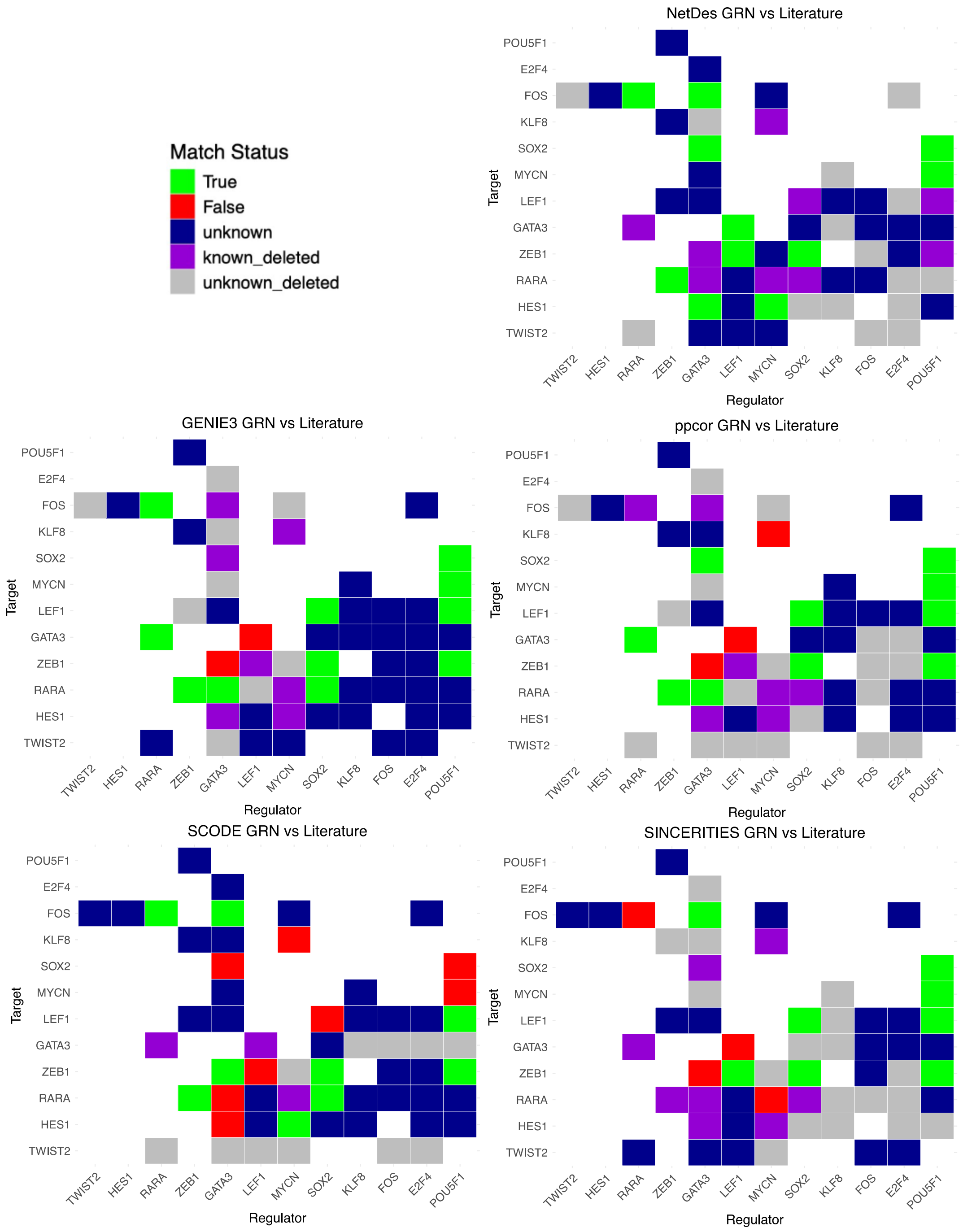

**Fig.S16. Comparison of inferred regulatory edges from each method against literature evidence.** This figure is related to **Fig.4E**. For each method, we illustrated the inferred GRN with the highest mapping percentage. Each panel shows a heatmap of regulator–target interactions for each inference method.

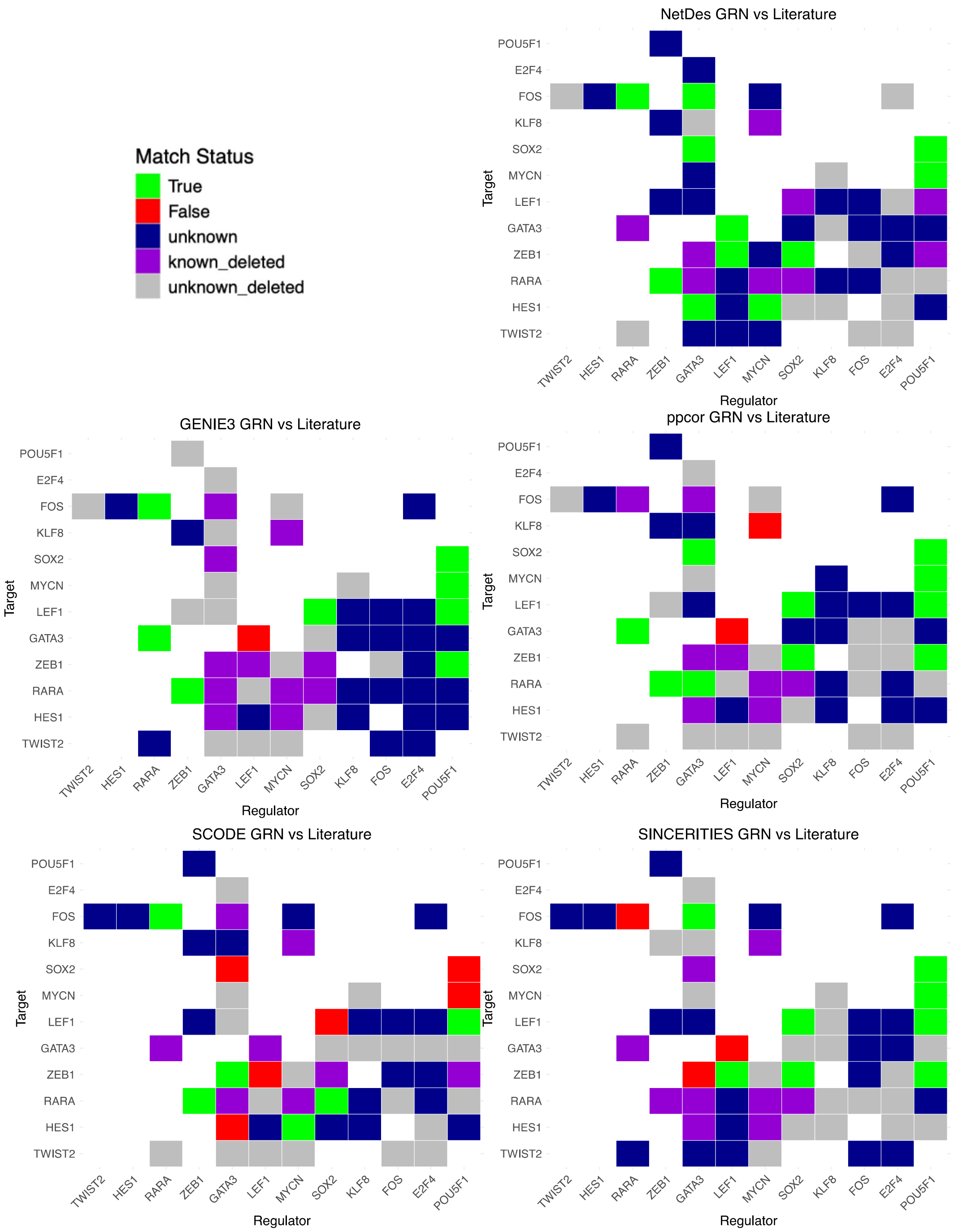

**Fig.S17. Comparison of inferred regulatory edges from each method against literature evidence.** This figure is related to **Fig.4E**. For each method, we illustrated the inferred GRN with the highest accuracy according to the literature comparison.

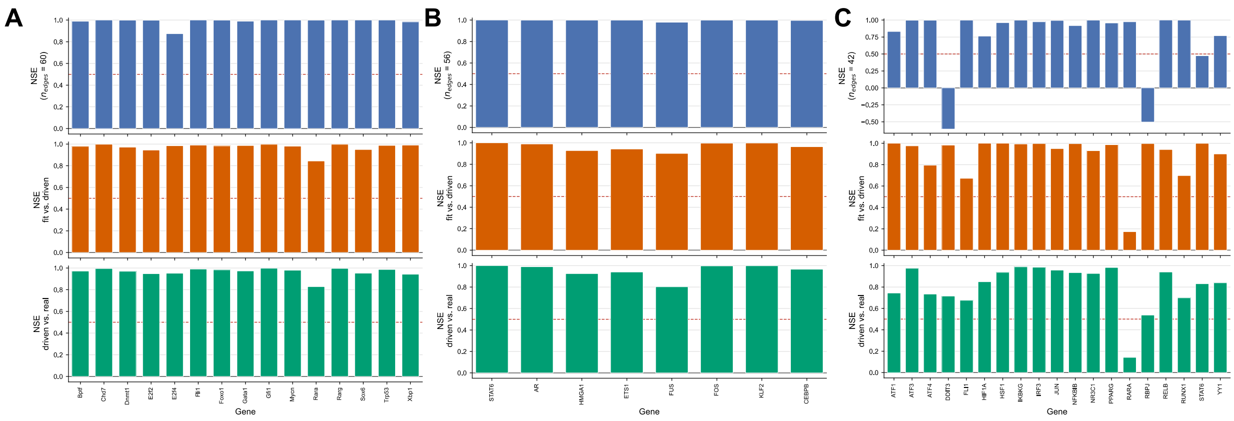

**Fig.S18. GRN quality check across the three cell-state-transition systems.** Bar plots of the Nash–Sutcliffe efficiency (NSE) between the simulated and reference gene expression trajectories, computed for each gene. **(A)** model fitting, **(B)** forward signaling, and **(C)** backward signaling. In each panel, the three rows (blue, orange, green) correspond to the EMT, ERY and DC datasets, respectively. A higher NSE (maximum of 1) indicates closer agreement between the simulated and observed trajectories.
